# Identifying gene function and module connections by the integration of multi-species expression compendia

**DOI:** 10.1101/649079

**Authors:** Hao Li, Daria Rukina, Fabrice P. A. David, Terytty Yang Li, Chang-Myung Oh, Arwen W. Gao, Elena Katsyuba, Maroun Bou Sleiman, Andrea Komljenovic, Qingyao Huang, Robert W. Williams, Marc Robinson-Rechavi, Kristina Schoonjans, Stephan Morgenthaler, Johan Auwerx

## Abstract

The functions of many eukaryotic genes are still poorly understood. We developed and validated a new method, termed GeneBridge, which is based on two linked approaches to impute gene function and bridge genes with biological processes. First, <u>G</u>ene-<u>M</u>odule <u>A</u>ssociation <u>D</u>etermination (G-MAD) allows the annotation of gene function. Second, <u>M</u>odule-<u>M</u>odule <u>A</u>ssociation <u>D</u>etermination (M-MAD) allows predicting connectivity among modules. We applied the GeneBridge tools to large-scale multi-species expression compendia—1,700 datasets with over 300,000 samples from human, mouse, rat, fly, worm, and yeast—collected in this study. Unlike most existing bioinformatics tools, GeneBridge exploits both positive and negative gene/module-module associations. We constructed association networks, such as those bridging mitochondria and proteasome, mitochondria and histone demethylation, as well as ribosomes and lipid biosynthesis. The GeneBridge tools together with the expression compendia are available at systems-genetics.org, to facilitate the identification of connections linking genes, modules, phenotypes, and diseases.

## Introduction

The identification of gene function and the integrated understanding of their roles in physiology are core aims of many biological and biomedical research projects — an effort that is still far from being complete (Edwards et al. 2011; Pandey et al. 2014; Dolgin 2017; Stoeger et al. 2018). Traditionally, gene function has been elucidated through experimental approaches, including the evaluation of the phenotypic consequences of gain- or loss-of-function (G/LOF) mutations (Austin et al. 2004; Dickinson et al. 2016), or by genetic linkage or association studies (Williams and Auwerx 2015). A large number of bioinformatics tools have been developed to predict gene function based on sequence homology (Marcotte et al. 1999; Radivojac et al. 2013; Jiang et al. 2016), protein structure (Roy et al. 2010; Radivojac et al. 2013; Jiang et al. 2016), phylogenetic profiles (Pellegrini et al. 1999; Tabach et al. 2013; Li et al. 2014), protein-protein interactions (Rolland et al. 2014; Hein et al. 2015; Huttlin et al. 2017), genetic interactions (Tong et al. 2004; Costanzo et al. 2010; Horlbeck et al. 2018), and co-expression (Langfelder and Horvath 2008; Warde-Farley et al. 2010; Greene et al. 2015; van Dam et al. 2015; Szklarczyk et al. 2016; Li et al. 2017; Obayashi et al. 2019).

With the development of transcriptome profiling technologies, thousands of high-throughput studies have generated a wealth of genome-wide data that has become a valuable resource for systems genetics analyses. A few web resources, including GEO (Barrett et al. 2013), ArrayExpress (Kolesnikov et al. 2015), GeneNetwork (Chesler et al. 2004), and Bgee (Bastian et al. 2008) amongst others, have created repositories of such expression data for curation, reuse, and integration. Several tools, such as GeneMANIA (Warde-Farley et al. 2010), GIANT (Greene et al. 2015), SEEK (Zhu et al. 2015), GeneFriends (van Dam et al. 2015), WeGET (Szklarczyk et al. 2016), COXPRESdb (Obayashi et al. 2019), WGCNA (Langfelder and Horvath 2008), and CLIC (Li et al. 2017), are able to assign putative new functions to genes by means of correlations or co-expression networks. At their core, these methods rely on the concept of guilt-by-association – that transcripts or proteins exhibiting similar expression patterns tend to be functionally related (Eisen et al. 1998). By using over-representation analyses on sub-networks or modules, one can then deduce aspects of gene functions.

However, these approaches generally depend on discrete sets of genes whose expression correlations exceed either a hard or soft threshold, which would strongly influence the final results. In addition, such analyses typically focus on positive or absolute values of correlations among datasets. The key polarity of interactions is often lost among gene products and linked modules (Warde-Farley et al. 2010; Greene et al. 2015; van Dam et al. 2015; Zhu et al. 2015; Li et al. 2017). Gene set analyses, such as gene set enrichment analysis (GSEA) (Subramanian et al. 2005), have been developed to identify processes or modules that are affected by certain genetic or environmental perturbations (Khatri et al. 2012). While GSEA removes the necessity of assigning a certain threshold, its application has mainly been limited to studying G/LOF models or environmental perturbations, where comparisons are inherently among discrete categories. This limits its applicability in most populations, in which variations among individuals are often subtle and continuous (Williams and Auwerx 2015).

Here we developed the GeneBridge toolkit that uses two interconnected approaches to improve upon the identification of gene function and to bridge genes to phenotypes using large-scale cross-species transcriptome compendia collected for this study. *First*, we describe a computational approach, named <u>G</u>ene-<u>M</u>odule <u>A</u>ssociation <u>D</u>etermination (G-MAD), to impute gene function. G-MAD considers expression as a continuous variable and identifies the associations between genes and modules. *Second*, we developed the <u>M</u>odule-<u>M</u>odule <u>A</u>ssociation <u>D</u>etermination (M-MAD) method to identify connections between modules based on the transcriptome compendia. The data and GeneBridge tools described here are available at systems-genetics.org, an open resource, which will facilitate the identification of novel connections between genes, modules, phenotypes, and diseases.

## Results

### Current status of gene annotations

Despite great efforts to annotate the cellular and physiological role of genes, many of their functions remain poorly understood. One of the most widely used resources of gene annotation for genes is the Gene Ontology (GO), which characterizes gene function based on three ontologies, i.e. biological process, molecular function, and cellular component (Ashburner et al. 2000). Over 54% (10,543 genes) of all the protein-coding genes in humans have no more than 10 annotations, including the uncurated IEA annotations (Inferred from electronic annotation) (Fig. 1A), whereas the most annotated gene *TP53* has more than 800 annotations (Fig. 1B). In fact, the top 20% most annotated genes have more than 64% of all annotations in GO (Fig. 1C). From these perspectives, it is clear that most human genes are still poorly annotated. The pattern is similar in other model species. Specifically, over 48% (10,166 genes) in mouse, 60% (11,833 genes) in rat, 61% (8,514 genes) in fly, 29% (5,885 genes) in worm, and 26% (1,566 genes) in yeast, have fewer than 10 entries in GO (Supplemental Fig. S1). This is also true for gene annotations retrieved from other sources, such as GeneRIF (Gene Reference Into Function) (Mitchell et al. 2003) (Fig. 1D-F), as well as for publications archived in PubMed (Barrett et al. 2013; Dolgin 2017) (Fig. 1G-I, Supplemental Fig. S1). The phenomenon that many genes are ignored in biological research has been pointed out before (Edwards et al. 2011; Pandey et al. 2014; Stoeger et al. 2018). Several possible reasons for this bias, such as prior knowledge, publication bias, and priorities of funding support have been raised (Edwards et al. 2011; Greene and Troyanskaya 2012; Stoeger et al. 2018). Therefore, an unbiased approach for gene function analysis would most likely provide many novel insights for future research.

**Figure 1.**
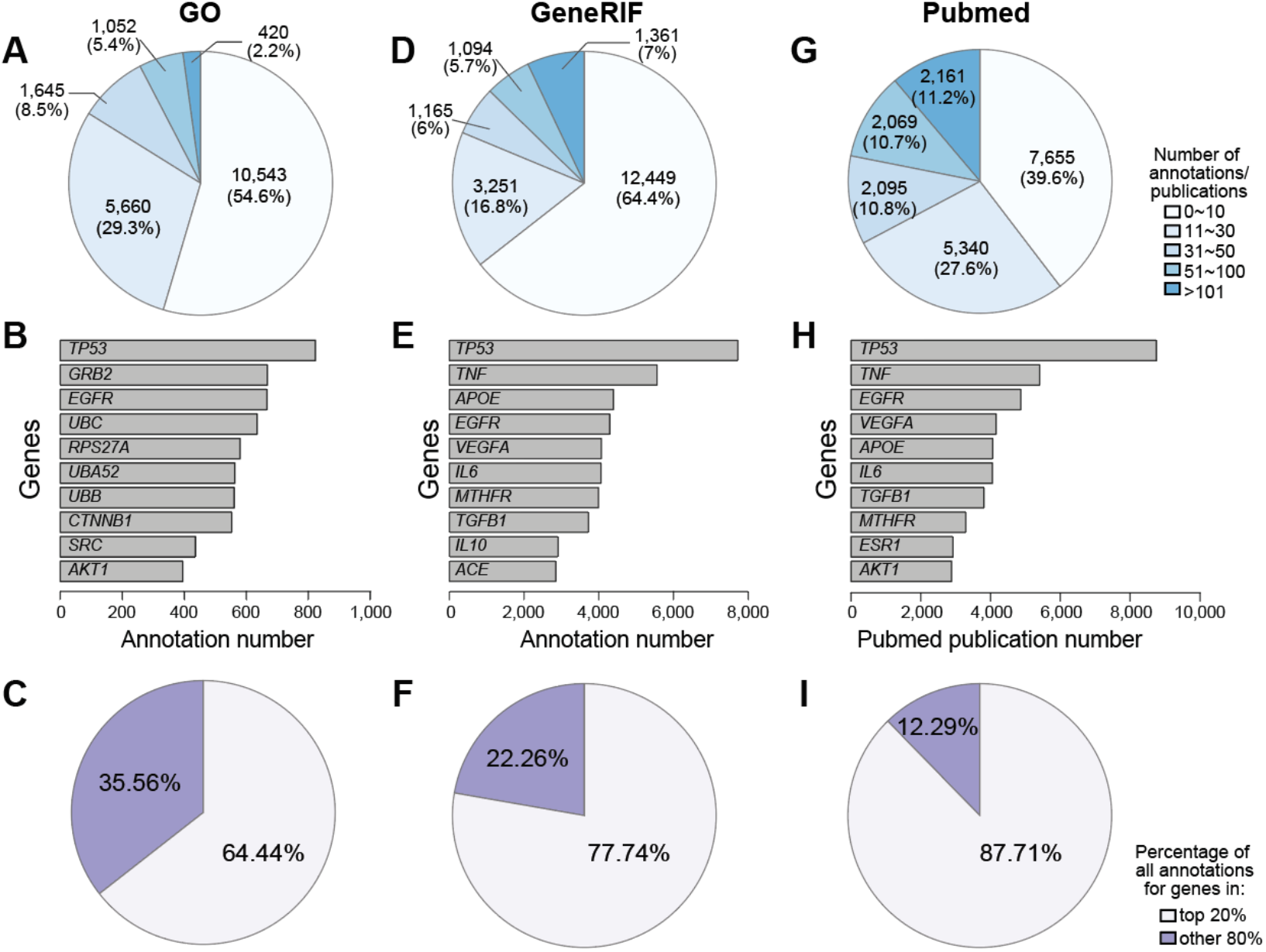
Known annotations for human genes. The number of annotations per gene for human genes in GO (**A**), GeneRIF (**D**), and number of publications in Pubmed (**G**). The top 10 genes with the most annotations/publications in GO (**B**), GeneRIF (**E**), and PubMed (**H**) are rank ordered. The percentage of all annotations/publications covering the top 20% most annotated genes in human (**C**, **F**, **I**).

### Gene-Module Association Determination (G-MAD)

Owing to the fact that a large number of genes are still not well annotated or even uncharacterized, we propose here a new computational strategy, “<u>G</u>ene-<u>M</u>odule <u>A</u>ssociation <u>D</u>etermination” (G-MAD), which uses expression data from large-scale cohorts to propose potential functions of genes. We use the term “modules” to refer the knowledge-based gene sets, ontology terms, and biological pathways from different resources for simplicity in the rest of the paper. The differences between gene sets or directed or undirected pathways are important in many contexts, but for our purpose they can be treated in the same manner as modules and will not be distinguished. The basic concept is similar to classic pathway/gene set analysis, i.e. genes that possess similar functions tend to have similar expression patterns (Subramanian et al. 2005). However, instead of using binary group settings (e.g., control *vs.* treatment, or wild-type *vs.* knockout) as commonly used in gene set analysis, we consider the continuous expression levels of the gene-of-interest across a population and determine its possible functions based on its co-expression patterns against all genes. We applied a competitive gene set testing method — Correlation Adjusted MEan RAnk gene set test (CAMERA), which adjusts for inter-gene correlations (Wu and Smyth 2012) — to compute the enrichment between gene-of-interest and biological modules. Gene-module connections with enrichment *p*-values that survived multiple testing corrections of the gene or module numbers were allocated connection scores of 1 or −1, based on the enrichment direction, and 0 otherwise. The results were then meta-analyzed across datasets, and gene-module association scores (GMAS) were computed as the averages of the connection scores weighted by the sample sizes and inter-gene correlation coefficients within modules (Fig. 2A).

**Figure 2.**
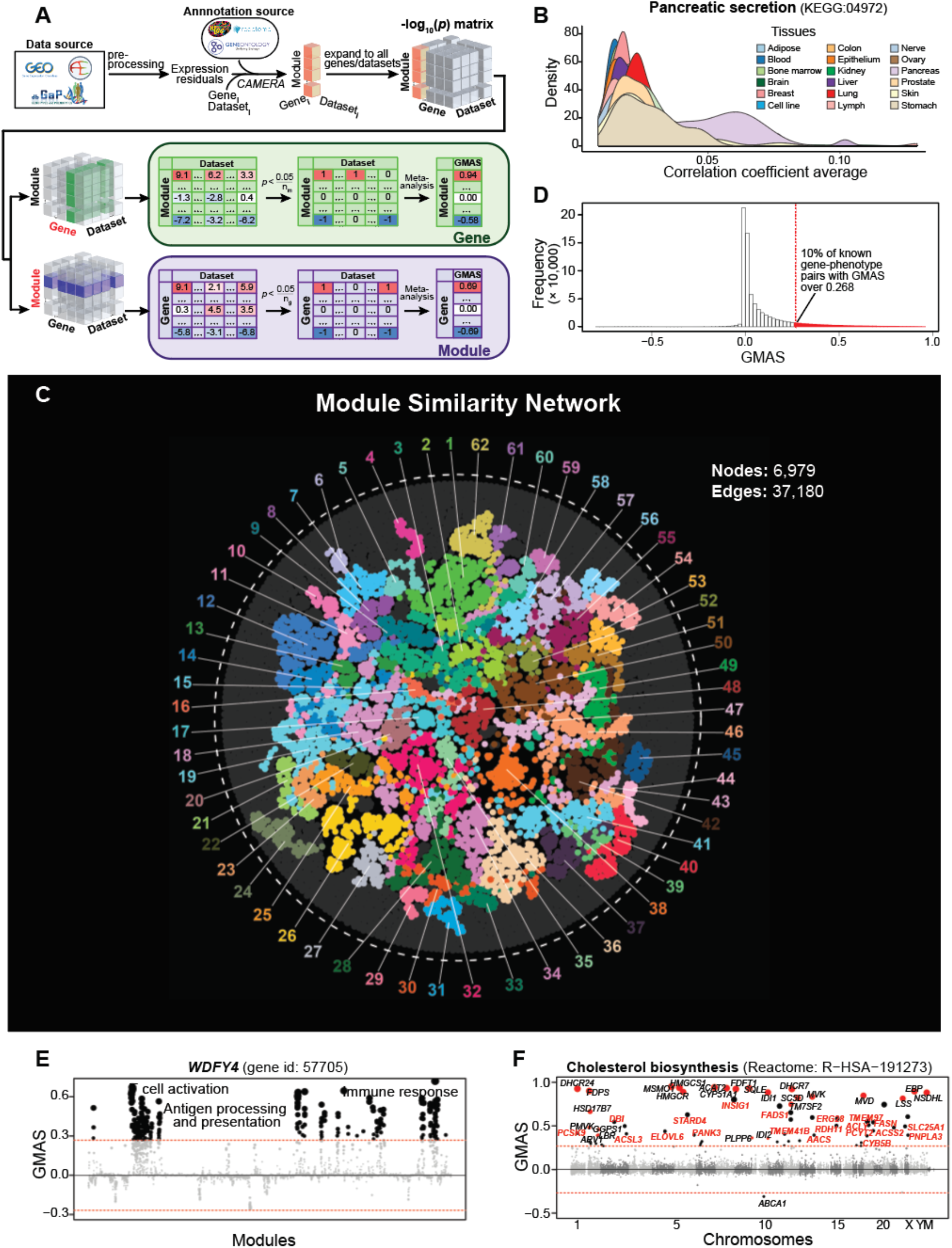
Gene-Module Association Determination (G-MAD). **A**, G-MAD methodology. See text and Materials and Methods for detailed description. **B**, Co-expressions among genes of pancreatic secretion module across tissues in human. The average correlation coefficient across all genes in the pancreatic secretion module in around 1,300 human expression datasets from 18 major tissues is used as to illustrate the co-expressions of this module across tissues. Genes in the pancreatic secretion module have higher co-expression in datasets from the pancreas compared to those from other tissues. **C**, Module similarity network showing the composition similarities across all module pairs. Modules were detected using community detection algorithm embedded in Gephi and indicated in different colors. The 10 most frequent words of the module terms in each module were used to represent the module, and can be found at Supplemental Table S2. **D**, Distribution of GMAS of all gene-module pairs with known connections in human. 10% of the known gene-module pairs have GMAS over 0.268. **E**, G-MAD revealed the potential role of *WDFY4* in T cell activation and immune response. The threshold of significant gene-module association is indicated by the red dashed line. Modules are organized by the module similarities. Known modules connected to *WDFY4* from annotations are shown in red dots (there is no known connected module for *WDFY4*), and other modules with GMAS over the threshold are shown in black dots. Dot sizes reflect the GMAS of *WDFY4* against the respective modules. **F**, G-MAD identified the involvement of known as well as 20 novel genes in cholesterol biosynthesis. The threshold of significant gene-module association is indicated by the red dashed line. Genes are organized by the genetic positions across chromosomes. Genes annotated to be involved in cholesterol biosynthesis are shown in red dots, and novel genes with GMAS over the threshold are shown in black dots. Novel genes conserved in human, mouse and rat are highlighted in red bold text.

We collected transcriptome datasets with over 80 samples from 6 species (human, mouse, rat, fly, worm and yeast), from GEO, ArrayExpress, dbGaP, GeneNetwork, and other data repository sources (Supplemental Table S1). For example, 1’337 datasets containing over 265’000 human samples with whole genome transcript levels were analyzed in this study (Supplemental Table S1). Genes annotated to some modules have higher co-expression in datasets from certain tissues than others (Supplemental Fig. S2A), suggesting the tissue-specific activation of these modules. For instance, genes involved in pancreatic secretion have much higher co-expressions in datasets obtained from pancreas (Fig. 2B). Genes belonging to “collecting duct acid secretion” module are highly co-expressed in kidney (Supplemental Fig. S2B-D), while genes in the “lamellar body” module are highly co-expressed in lung (Supplemental Fig. S2E-G).

One should be aware of the fact that modules can overlap partially or completely. For example, GO categories have a hierarchical structure. Each GO term has "parent" terms (related usually by "part of" or "is a" relations), and all genes annotated to the term will also be annotated to its parents. Symmetrically, a GO term can be the parent term of other GO terms (Ashburner et al. 2000). In addition, modules from different sources can be very similar in composition. For example, oxidative phosphorylation (84 genes) from GO biological process, and respiratory chain (80 genes) from GO cellular component have 65 genes (66%) in common in humans. Therefore, we computed the similarities across all modules, and generated a global module similarity network. As expected, redundant modules formed clusters in the network, and we were able to extract 62 distinct module clusters in the human module similarity network (Fig. 2C, Supplemental Table S2).

We assessed the performance of G-MAD in prioritizing known genes for modules through cross validations. We then compared the area under the receiver operating characteristic (ROC) curve (AUC) with the ones obtained from WeGET, a method predicting novel genes for various modules based on weighted co-expression of around 1,000 expression datasets (Szklarczyk et al. 2016). G-MAD exhibits better predictive performance than WeGET (Supplemental Fig. S3). Furthermore, in order to determine the threshold of significance of gene-module associations, we computed the GMAS of all the known gene-module pairs. To be stringent in proposing novel gene-module associations, we consider only 10% of all the known gene-module pairs as significant, and picked a GMAS threshold of 0.268 (Fig. 2D). With this threshold, we saw only 0.24% of unknown gene-module pairs are significant, which is 40 times less than the known pairs.

The gene-module connections predicted by G-MAD provide a resource, which researchers can use as a reference when annotating gene functions. We describe below some examples on how the G-MAD results can be used to facilitate the discovery of novel gene functions or the identification of new members of modules. *WDFY4* was recently annotated as a crucial gene in activating immunological T cells in antiviral and antitumor immunity through a functional CRISPR screen (Theisen et al. 2018). Through G-MAD, we found that *WDFY4* is indeed associated with antigen processing, T cell activation, and immune response in human, mouse, and rat (Fig. 2E, Supplemental Fig. S4A-B), verifying its functions conserved across species. Cholesterol is critical in cell differentiation and growth. We identified 20 genes (*AACS*, *ACLY*, *ACSL3*, *ACSS2*, *CYB5B*, *DBI*, *ELOVL6*, *ERG28*, *FADS1*, *FASN*, *INSIG1*, *PANK3*, *PCSK9*, *PCYT2*, *PNPLA3*, *RDH11*, *SLC25A1*, *STARD4*, *TMEM41B*, *TMEM97)* associated to cholesterol biosynthesis conserved in human, mouse, and rat (Fig. 2F, Supplemental Fig. S4C-E). Several of these genes, including *FASN* (Carroll et al. 2018) and *TMEM97* (Bartz et al. 2009), have already been described to have relevant functions in cholesterol metabolism.

G-MAD can also highlight tissue-specific gene-module associations using datasets from specific tissues. *EHHADH* is a peroxisomal protein highly expressed in liver and kidney (Fig. 3A) (Uhlen et al. 2015). Although best known for its key role in the peroxisomal oxidation pathway, recent report demonstrated that *EHHADH* mutations cause renal Fanconi’s syndrome (Klootwijk et al. 2014). G-MAD of *EHHADH* in liver and kidney identifies its conserved role in peroxisome and fatty acid oxidation, and also recovers its specific functions in liver (e.g. bile acid biosynthesis) and kidney (e.g. brush border membrane) (Fig. 3B-E, Supplemental Table S3). *SLC6A1* is one of the major gamma-aminobutyric acid (GABA) transporters in the neurotransmitter release cycle in brain (Carvill et al. 2015). However, *SLC6A1* is also highly expressed in the liver (Supplemental Fig. S5A), and its function in liver remains poorly understood. G-MAD of *SLC6A1* in all datasets and only datasets from brain confirms its function as neurotransmitter transporters in GABA release cycle (Supplemental Fig. S5B-C), while G-MAD using datasets from liver identifies its possible role in carboxylic acid transport and metabolism (Supplemental Fig. S5D-E, Supplemental Table S4).

**Figure 3.**
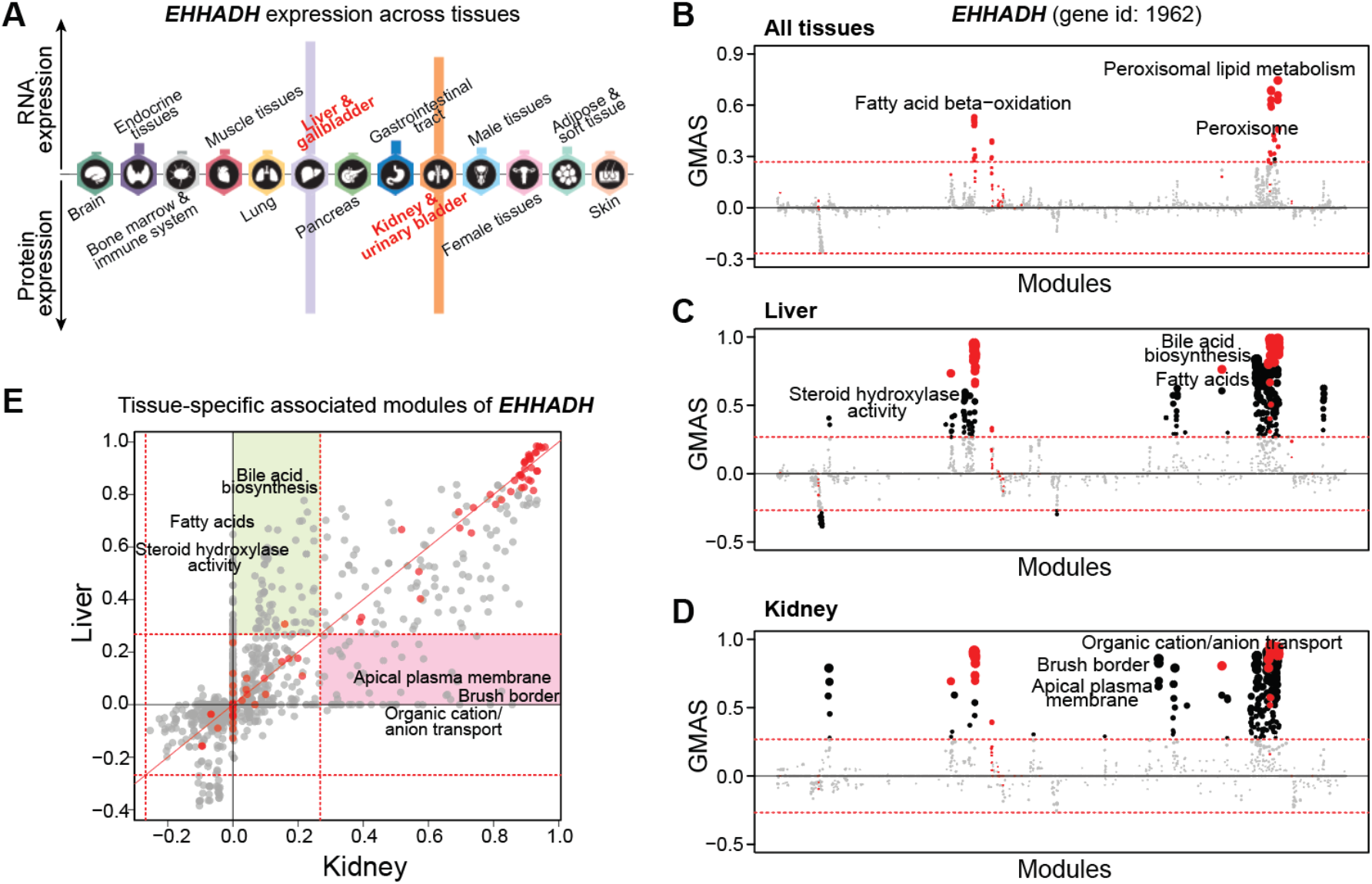
G-MAD identifies tissue-specific associated modules for *EHHADH* by using datasets from different tissues. **A**, Expression patterns of *EHHADH* across tissues. The figure was adapted from the Human Protein Atlas. **B**-**D**, G-MAD of *EHHADH* in human using datasets from all tissues (**B**), from liver (**C**), or from kidney (**D**). The threshold of significant gene-module association is indicated by the red dashed line. Modules are organized by their similarities. Known modules connected to *EHHADH* from gene annotations are shown in red dots, and other modules with GMAS over the threshold are shown by black dots. **E**, Comparison of G-MAD results of *EHHADH* in liver and kidney. Known modules connected to *EHHADH* are shown in red dots. The threshold of significant gene-module association is indicated by the red dashed line. Modules significantly associated with *EHHADH* only in one specific tissue are highlighted. The comparison of the association results of *EHHADH* in liver and kidney can be found at Supplemental Table S3.

### G-MAD determines novel genes linked to mitochondria

Mitochondria are the main powerhouses of cells and harvest energy in the form of ATP through mitochondrial respiration. There are around 1,100 genes known to encode mitochondria-localized proteins (mito-proteins), depending on the source used (e.g. 1,158 mito-proteins in Mitocarta (Calvo et al. 2016), 1,074 in Human Protein Atlas (Uhlen et al. 2015)); however, many of these genes remain uncharacterized, and the list of mito-proteins is still incomplete (Williams et al. 2018).

By using the genes annotated to be involved in respiratory electron transport chain (ETC, Reactome: R-HSA-611105), we searched for genes potentially related to respiratory electron transport, by applying G-MAD to expression datasets in human, mouse, and rat. As expected, genes annotated in the ETC module are strongly enriched; moreover, other known ETC genes that were not included in the module were also positively enriched, providing proof that G-MAD can recover known gene functions (Fig. 4A, Supplemental Fig. S6A-B). Based on G-MAD results from human, mouse and rat, there were 707 genes showing conserved associations with the ETC (Fig. 4B). Many of these genes, for example *DMAC1*/*C9orf123 (Arroyo et al. 2016; Stroud et al. 2016; Horlbeck et al. 2018)*, *NDUFAF8*/*C17orf89 (Floyd et al. 2016)*, and *FMC1*/*C7orf55* (Lefebvre-Legendre et al. 2001; Li et al. 2017) were not included in the respiratory electron transport module, but have been recently validated to be involved in mitochondrial respiration (Fig. 4B, Supplemental Table S5). *DDT* is among the top genes associated with the ETC (Fig. 4A-B), and there is no previous study linking it to mitochondria. G-MAD reveals that *DDT* is strongly associated with mitochondrial respiration across different species, including the invertebrate *C. elegans* (Fig. 4C-D, Supplemental Fig. S6C-G), suggesting a conserved role of *DDT* in mitochondria. We experimentally validated this finding through RNAi-mediated *DDT* knockdown in HEK293 cells, which led to reduced transcript levels of genes encoding for the ETC subunits (Fig. 4E) and decreased oxygen consumption rate (OCR) (Fig. 4F, Supplemental Fig. S5H), confirming that *DDT* impacts mitochondrial respiration. Similarly, we also confirmed the involvement of *BOLA3* in the ETC using G-MAD and experimental validations (Cameron et al. 2011) (Supplemental Fig. S7).

**Figure 4.**
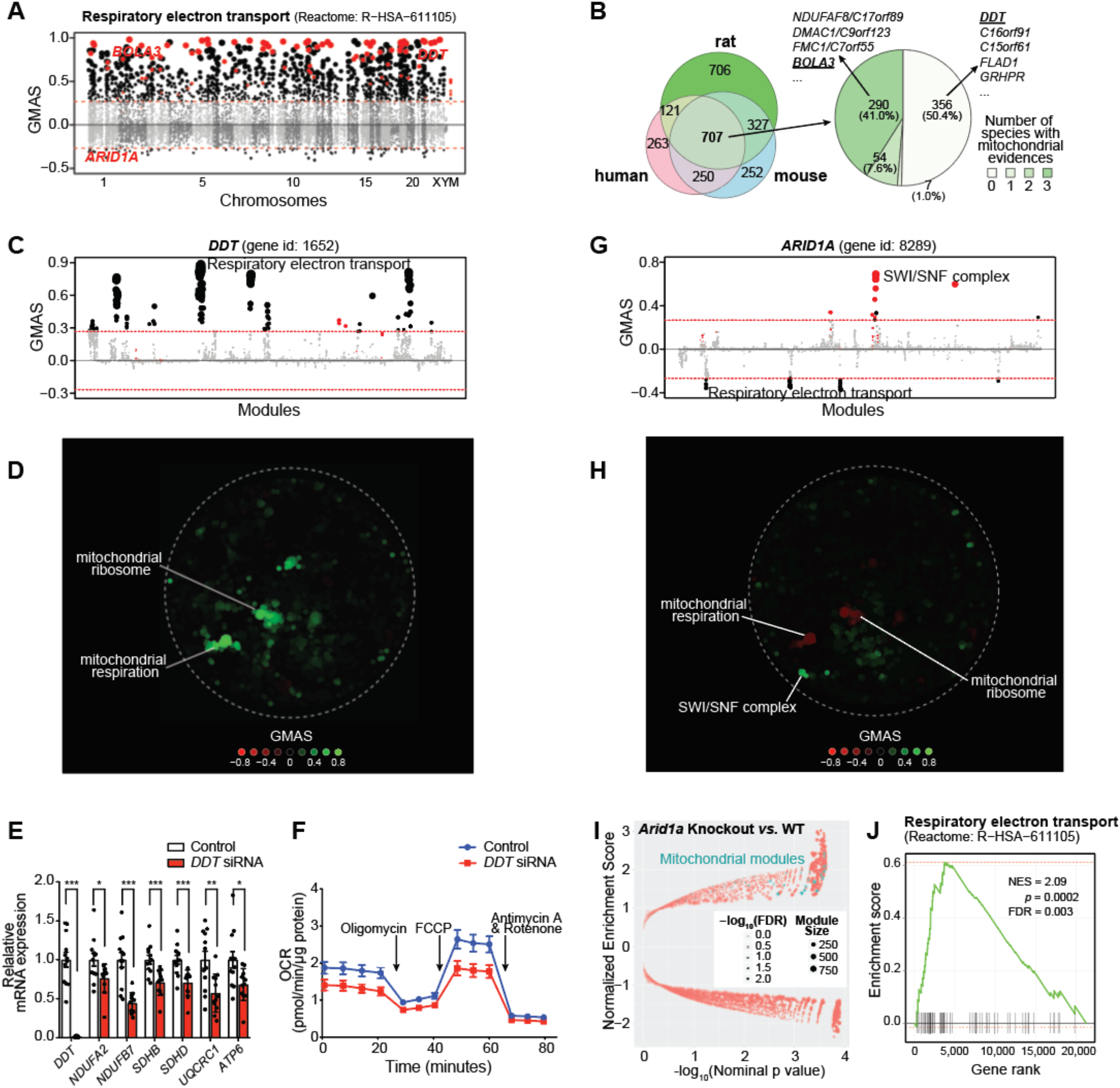
G-MAD predicts novel genes linked to mitochondria. **A**, G-MAD Manhattan plot of the respiratory electron transport (Reactome: R-HSA-611105) module in human. Genes are arranged based on their genetic positions, and genes annotated to be involved in the module are colored red. Genes with absolute GMAS over 0.268 are considered significantly associated. *DDT*, *BOLA3*, and *ARID1A* are labeled. **B**, Venn diagram of novel genes associated with respiratory electron transport module in human, mouse and rat. 707 genes were predicted to be mito-proteins by G-MAD in all three species. 351 genes, including *DMAC1*/*C9orf123*, *NDUFAF8*/*C17orf89*, *FMC1*/*C7orf55*, and *BOLA3*, were recently annotated to be involved in mitochondrial respiration in at least one species. While 356 genes, including *DDT*, *C16orf91*, *C15orf61*, *FLAD1*, and *GRHPR*, have not been previously linked with mitochondria based on the current annotations. The association results for all genes in human, mouse and rat can be found at Supplemental Table S4. **C**, *DDT* associates with mitochondrial respiratory chain modules in human. The threshold of significant gene-module association is indicated by the red dashed line. Modules are organized by module similarities. Known modules connected to *DDT* from annotations are highlighted in red, and other modules with GMAS over the threshold are colored in black. Dot sizes reflect the GMAS of *DDT* against the respective modules. **D**, Module similarity network showing the modules associated with *DDT*. Modules are plotted based on their layout in Fig. **2C** and colored based on their GMAS against *DDT*. **E**, Silencing *DDT* expression in HEK293 cells decreases expression levels of indicated genes involved in mitochondrial respiratory chain complexes. Error bars represent standard errors. *, *p* < 0.05; **, *p* < 0.01; ***, *p* < 0.001. n=12. **F**, *DDT* knockdown leads to the reduction of oxygen consumption rate (OCR) as a reflection of mitochondrial respiration in human HEK293 cells. Addition of specific mitochondrial inhibitors, including the oligomycin (ATPase inhibitor), FCCP (uncoupling agent), and rotenone/antimycin A (electron transport chain inhibitors) are indicated by arrows. **G**, *ARID1A* negatively associates with mitochondrial respiratory chain in human. The threshold of significant gene-module association is indicated by the red dashed line. Modules are organized by the module similarities. Known modules connected to *ARID1A* from extant annotations are highlighted in red, and other modules with GMAS over the threshold are colored in black. Dot sizes are proportional to GMAS of the respective modules. **H**, Module similarity network showing the modules associated with *ARID1A*. Modules are colored based on their GMAS against *ARID1A*. **I**, *Arid1a* uterine-specific knockout mice showed positive enrichment in mitochondrial respiration modules. Nominal *p*-values from the GSEA results are used to plot against normalized enrichment score (NES), with dot sizes indicating the number of genes in the modules and transparencies indicating the false discovery rate (FDR). **J**, Enrichment plot showing the enrichment of genes included in respiratory electron transport in uterus-specific *Arid1a* knockout mice compared to wild-type controls. Genes are ranked based on the fold change between *Arid1a* knockout and wild-type mice, and the ranking positions of genes in respiratory electron transport are labeled as vertical black bars. NES, normalized enrichment score. FDR, false discovery rate.

Contrary to the existing methods that predict only positive gene-module associations based on gene co-expression, G-MAD is also able to exploit negative associations. For example, *ARID1A* exhibits significant negative associations with the respiratory electron transport in human and mouse (Fig. 4A,G-H, Supplemental Fig. S8). *ARID1A* is a known member of the SWI/SNF family, and the inactivating mutations of SWI/SNF complex genes (mainly *SMARCA4* and *ARID1A*) have recently been linked to increased expression of ETC genes and mitochondrial respiration (Lissanu Deribe et al. 2018). To further validate its regulatory role, we checked an extant public dataset from mice with uterus-specific *Arid1a* knock-out (Kim et al. 2015), and confirmed that dysfunction of *Arid1a* led to the increased expression of mitochondrial genes (Fig. 4I), especially those involved in respiratory electron transport (Fig. 4J).

### Module-Module Association Determination (M-MAD)

Biological processes and modules, such as metabolism, cellular signaling, biogenesis, and degradation are interconnected and coordinated (Barabasi et al. 2011). However, there are few reports exploring the connections between modules in a systematic fashion (Li et al. 2008). Here we extend G-MAD to develop <u>M</u>odule-<u>M</u>odule <u>A</u>ssociation <u>D</u>etermination (M-MAD) to investigate the connections between modules based on the expression compendia. Results for individual modules against all genes, obtained from G-MAD, were used to compute their associations against all modules. The enrichment scores of all genes for the target module were used as the gene-level statistics to calculate the enrichment against all modules using CAMERA (Wu and Smyth 2012). The resulting enrichment *p*-values across modules were transformed to 1, 0, or –1 based on the Bonferroni threshold, and then meta-analyzed across all datasets to obtain the module-module association scores (MMAS) (Fig. 5A).

**Figure 5.**
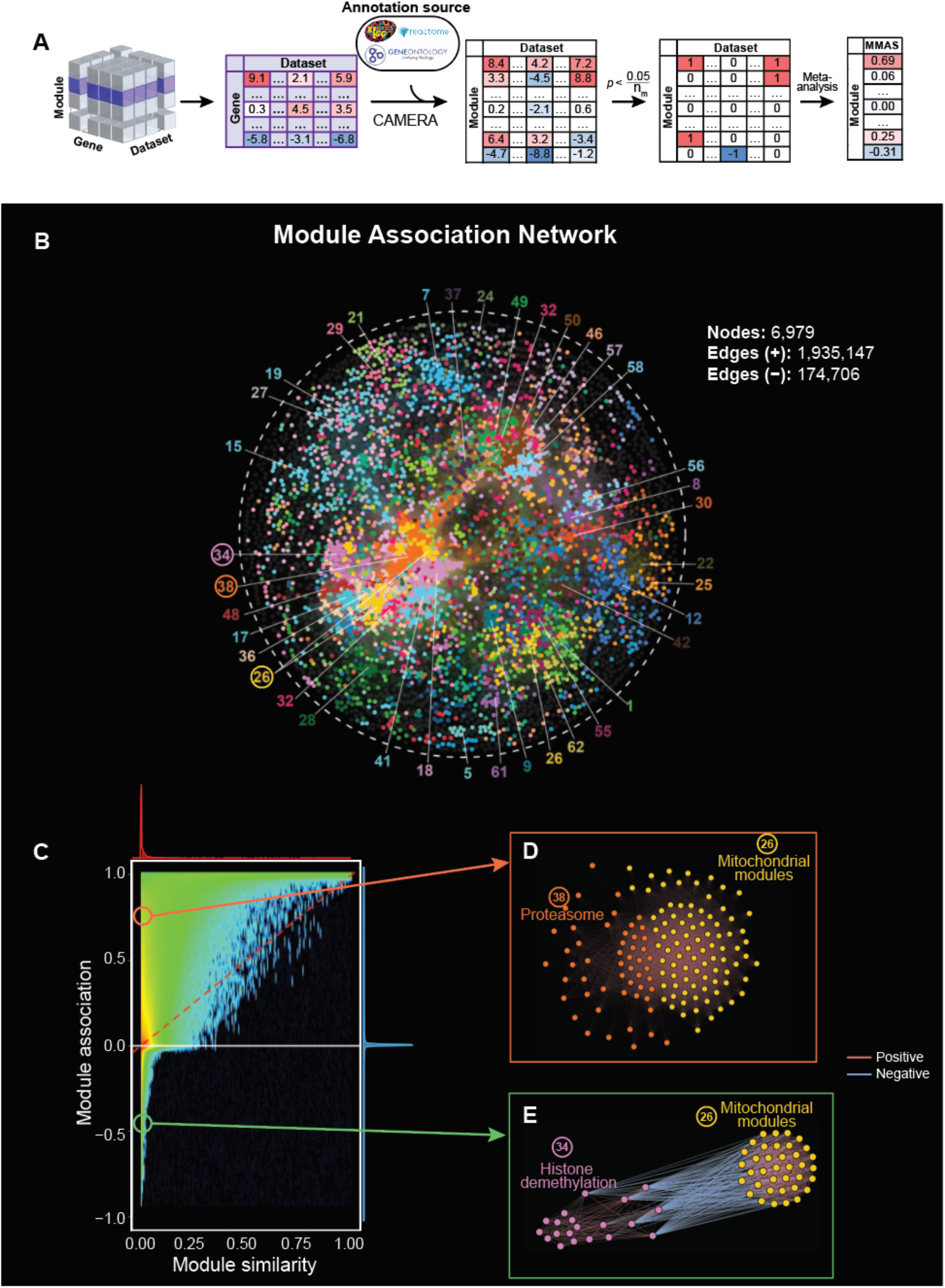
Module-Module Association Determination (M-MAD) reveals module connections. **A**, Scheme of the M-MAD methodology in detecting module connections. Intermediate results of G-MAD for all modules are further processed and used as the basis of M-MAD. The -log_10_(*p*) values of G-MAD for the target module against all genes in each dataset are used as the gene statistic for the module, and connections between the target module and all modules are calculated using CAMERA. The results are then meta-analyzed by taking the sample sizes and inter-gene correlations of all datasets to compute the <u>m</u>odule-<u>m</u>odule <u>a</u>ssociation <u>s</u>core (MMAS) between modules. **B**, Module association network showing the connections across all modules. Colors of nodes represent the module modules defined in the global module similarity network in Fig. **2C**. Module clusters with respective colors are identified and labeled. Modules used as examples in following figures are highlighted in circle. **C**, Comparison of pairwise module connections derived from module similarities in Fig. **2D** and associations (from M-MAD) in Fig. **5B**. A red dashed line is plotted when the pairwise module similarity equals association. The distributions of module similarity and association scores are illustrated in the top and at the right of the plot and are colored in red and blue, respectively. Two examples of novel module connections are labeled. **D**-**E**, Subnetworks showing the association between mitochondrial and proteasomal modules (**D**), and mitochondrial and histone demethylation modules (**E**). Edges colors indicate the significance of module connections, with red as positive and blue as negative.

Module-module associations with an absolute MMAS of over 0.268, corresponding to 4% of the total number of module pairs, were considered significant and were used to construct a module association network (Fig. 5B). Modules were represented as nodes with the same colors as the module clusters from Fig. 2C. While the module <u>*similarity*</u> network in Fig. 2C is based solely on existing gene annotations, the module <u>*association*</u> network relies on analyzing the full expression datasets. It can thus reveal new biological connections among modules, which were not included in literature-based annotations. We compared the two networks (Supplemental Fig. S9) obtained from module similarity (Fig. 2C) and module association (Fig. 5B). Interestingly, there are numerous module pairs with no similarity/overlap of annotated genes, but with high association based on expression (M-MAD) (Fig. 5C). Moreover, many module pairs have predicted negative associations (Fig. 5C). Therefore, these results provide a resource for hypothesis generation and validation of the module connections.

By applying M-MAD, we observed a strong positive link between mitochondrial modules and the proteasome (Fig. 5D, Supplemental Fig. S10A-C). Most of the genes encoding proteasome subunits exhibit remarkable association with the ETC in human and mouse (Supplemental Fig. S10G), indicating a conserved co-regulatory mechanism. Dysfunction of mitochondria and the ubiquitin-proteasome system (UPS) are hallmarks of aging and aging-related neurodegenerative diseases, such as Alzheimer’s, Parkinson’s, and Huntington’s diseases (Ortega and Lucas 2014; Ross et al. 2015; D’Amico et al. 2017). Abnormalities that perturb the crosstalk between these two modules have been demonstrated to contribute to the pathogenesis of these diseases and several mechanisms have been proposed (D’Amico et al. 2017; Harrigan et al. 2017). It has also been shown that ETC disruption leads to proteasome impairment (D’Amico et al. 2017), while conversely the inhibition of the UPS causes mitochondrial dysfunction (Ross et al. 2015).

Similar to G-MAD, M-MAD can also predict negative connections between modules. For example, we found strong negative connections between histone demethylation processes and mitochondrial modules (Fig. 5E, Supplemental Fig. S10D-F). The link between epigenetics and mitochondria is a research focus for many groups, including ours (Schroeder et al. 2013; Merkwirth et al. 2016; Tian et al. 2016). It has been reported that mitochondrial dysfunction affects histone methylation, and conversely histone lysine demethylases can impact mitochondrial functions (Merkwirth et al. 2016). Most of the histone lysine demethylases showed negative associations with the ETC in human and mouse (Supplemental Fig. S10G), suggesting a conserved negative connection between histone demethylation and mitochondrial function.

As another example of M-MAD, we investigated modules connected with lipid biosynthetic modules. Interestingly, ribosome modules exhibited strong negative association with lipid biosynthetic modules (Fig. 6A-B, Supplemental Fig. S11A-B). This is in line with our previous finding that a ribosomal protein, *Rpl26*, negatively correlates with body weight and fat mass (Li et al. 2018). In support of this connection, liver and adipose transcripts of most ribosomal protein genes negatively correlated with metabolic phenotypes, such as body weight, fat mass, and cholesterol levels in the BXD mouse cohort (Wu et al. 2014) (Fig. 6C, Supplemental Fig. S11C), as well as in a CAST/EiJ and C57BL/6J F2 intercross (Schadt et al. 2008) (Fig. 6D, Supplemental Fig. S11D). Finally, RNAi targeting 9 of the identified ribosomal protein genes out of total 13 tested led to the accumulation of lipid droplets in *C. elegans* (Fig. 6E, Supplemental Fig. S11E-G), further validating the robustness of the lipid synthesis-ribosome connection across species.

**Figure 6.**
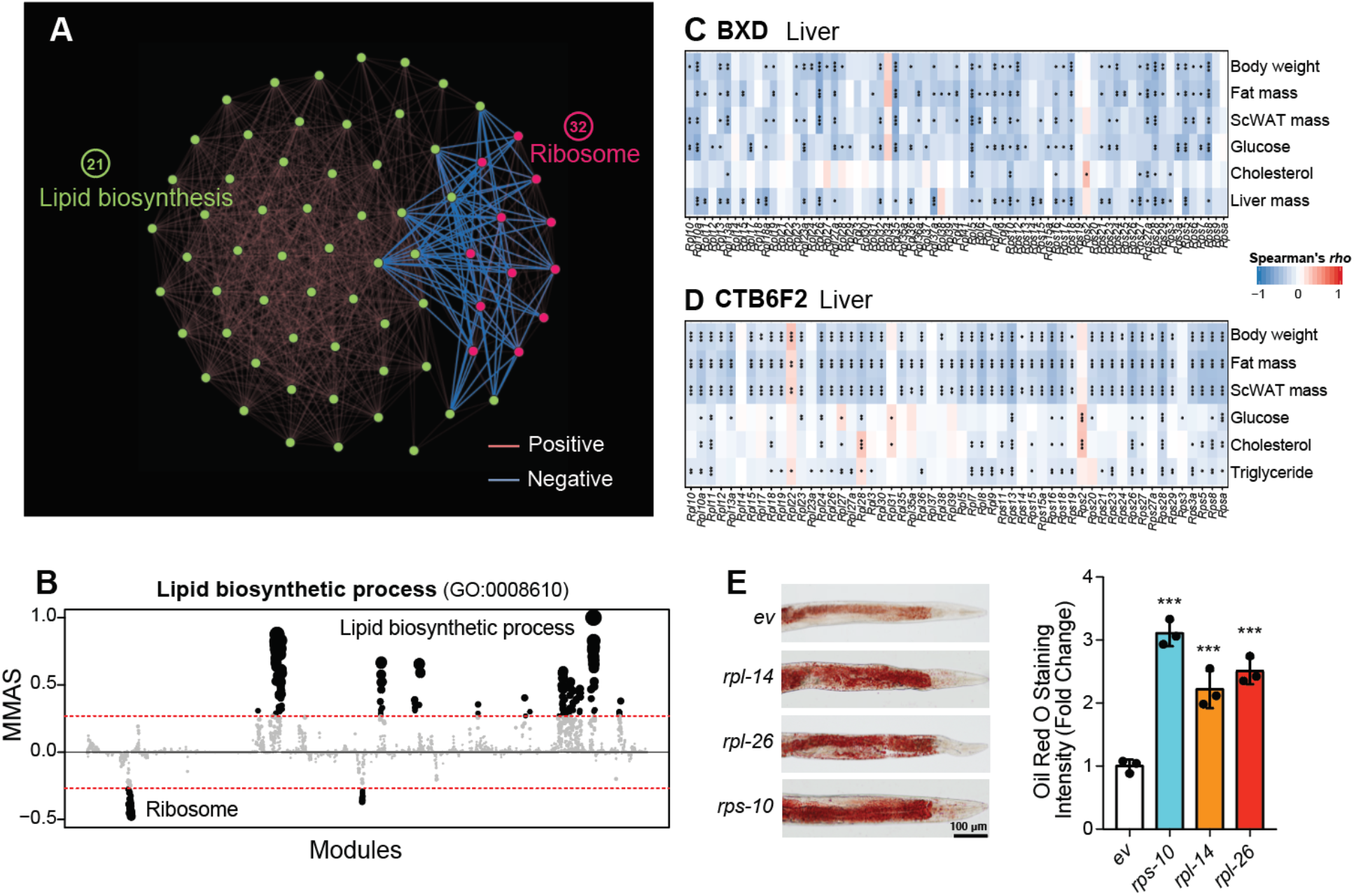
M-MAD reveals a negative association between the ribosome and lipid biosynthetic modules. **A**, Subnetwork for the ribosome and lipid biosynthetic modules. The colors of the edges indicate the significance of module connections, with red as positive and blue as negative. **B**, Lipid biosynthetic process negatively connected with ribosomal modules in human. The threshold of significant module-module connection is indicated by the red dashed line. Modules are organized by the module similarities. Dot sizes are proportional to MMAS of the respective modules. **C**-**D**, Transcripts of genes encoding for ribosomal proteins in the liver negatively correlate with metabolic traits, such as body weight, fat mass, plasma glucose and cholesterol levels, in the BXD (**C**) and CTB6F2 (**D**) mouse cohorts. *, *p* < 0.05; **, *p* < 0.01; ***, *p* < 0.001. **E**, Feeding adult *C. elegans* with RNAi clones of ribosomal proteins, including *rps-10*, *rpl-14*, and *rpl-26*, results in the accumulation of lipids, as reflected by Oil Red O staining. Experimental scheme and additional examples are shown in Supplemental Fig. S7. ***, *p* < 0.001. *ev*, empty vector. n=3.

## Discussion

Significant efforts in biological research have been devoted to defining the molecular and physiological functions of genes. However, many genes are still not well annotated and even remain uncharacterized (Edwards et al. 2011; Dolgin 2017; Stoeger et al. 2018). Here we developed an approach, termed G-MAD, to facilitate the identification of novel gene functions and to establish robust connections between genes and modules. Using transcriptome datasets from cohorts ranging from human to mouse, rat, fly, worm, and yeast, we identified millions of gene-module connections, many of which are novel. Unlike usual co-expression analyses for predicting gene functions, G-MAD can identify not only positive gene-module connections, but also negative associations between genes and modules or processes. We illustrated the predictive power of G-MAD in revealing potential gene-module connections using the mitochondrial electron transport chain (ETC) module as an example. 707 genes were consistently associated with the ETC in human, mouse and rat, of which *DDT* and *BOLA3* were validated through experiments. A negative connection between *ARID1A*, a member of the SWI/SNF family, and the ETC was also identified using G-MAD, which was consistent with a report that inactivation of SWI/SNF complex increased mitochondrial respiration (Lissanu Deribe et al. 2018). Meanwhile, tissue-specific functions of genes, for example *EHHADH* and *SLC6A1*, can also be identified using datasets derived from respective tissues.

In addition, we extended G-MAD to M-MAD, to uncover connections between modules. Association scores of one module against all genes from G-MAD were used to compute its associations with all modules. Similar to G-MAD, M-MAD can identify both positive and negative module associations. For example, in humans we identified around 2,000,000 associations between all modules, over 170,000 of which negative. We constructed a module association network based on these connected modules, and compared it to the module similarity network. Interestingly, many of the associated module pairs have low or no similarities in gene compositions. By applying M-MAD on the ETC module, we discovered a conserved connection between mitochondria and the proteasome in various organisms (D’Amico et al. 2017). In addition, we identified negative associations between histone lysine demethylation and mitochondrial modules, underscoring the inverse connection between epigenetic regulation and mitochondrial function (Schroeder et al. 2013; Merkwirth et al. 2016; Tian et al. 2016). Moreover, we discovered and validated a novel negative regulatory role of ribosomal proteins on lipid biosynthesis (Li et al. 2018).

In summary, we described here a set of approaches to identify gene function and module connectivity, that we collectively termed GeneBridge, to reflect their capacity to bridge genes to biological functions and phenotypes. The GeneBridge toolset is accessible through our open web resource (systems-genetics.org) to the research community for hypothesis generation or validation. It should be noted that although only protein-coding genes were included in our analysis, the same approach can be applied to non-coding genes to reveal their potential functions. Similarly, GeneBridge can also be utilized to identify novel gene-disease associations based on known disease-associated genes from databases, such as the Human Disease Ontology (DO) (Schriml et al. 2019) or DisGeNET (Pinero et al. 2017). The GeneBridge toolkit could also be applied to large-scale proteomics datasets after correcting for the background of all measured proteins. Integration of GeneBridge with other well-established databases, such as BioGRID (Stark et al. 2006) and STRING (Szklarczyk et al. 2015), will facilitate the investigation of the connections between genes, modules, and diseases.

## Methods

### Gene annotations / Modules

Gene ontology (GO) annotations (Ashburner et al. 2000) were downloaded from http://www.geneontology.org/ on Oct 4, 2017, with versions indicated by submission date below. Gene Reference Into Function (GeneRIF) (Mitchell et al. 2003) was downloaded from ftp://ftp.ncbi.nih.gov/gene/GeneRIF/ on Oct 11, 2017. Publication information from PubMed was downloaded from ftp://ftp.ncbi.nlm.nih.gov/gene/DATA/gene2pubmed.gz on Mar 15, 2018.

Module data for all the species were retrieved from GO (Ashburner et al. 2000), Kyoto Encyclopedia of Genes and Genomes (KEGG) (Kanehisa et al. 2012), and Reactome (Croft et al. 2011). Annotations from GO with evidence codes of IEA (inferred from electronic annotation), ND (No biological data available), NR (Not recorded), NAS (Non-traceable author statement) were removed from the analysis. The parent-child hierarchical structure of GO was ignored. All modules, including the redundant modules (modules with similar gene components), as well as parent-child modules, were considered as independent in the analysis.

Modules with less than 15 genes or larger than 1,000 genes were excluded, resulting in 6,979, 7,489, 7,462, 3,811, 2,495, and 2,381 modules for human, mouse, rat, fly, worm, and yeast, respectively, for the analysis.

### Module similarity calculation

Similarity between two modules were defined as the Jaccard index 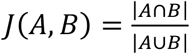, i.e. the number of genes in A and B divided by the number of genes in A or B. It measures the intersection between the modules as a fraction of the total size.

### Gene expression across tissues

Expression patterns of *EHHADH* and *SLC6A1* in mRNA and protein levels across human tissues were obtained from the Human Protein Atlas (Uhlen et al. 2015), and are available from v18.proteinatlas.org/ENSG00000113790-EHHADH/tissue and v18.proteinatlas.org/ENSG00000157103-SLC6A1/tissue, respectively.

### Transcriptome datasets

Human GTEx transcriptome datasets were downloaded from https://www.gtexportal.org (GTEx_Consortium 2013). Most of the microarray and RNAseq datasets were downloaded from GEO (Barrett et al. 2013) and ArrayExpress (Kolesnikov et al. 2015), with processed human and mouse RNAseq datasets obtained from ARCHS4 (Lachmann et al. 2018). The rest of the datasets were downloaded from other sources, including the database of Genotypes and Phenotypes (dbGaP) (Mailman et al. 2007), Mouse phenome database(Bogue et al. 2018), and other data repository websites. Data from single cell RNA-seq were excluded from the study because they contain the many zero counts. Detailed information can be found at systems-genetics.org/datasets.

### Data preprocessing of transcriptome datasets

For microarray datasets, the expression for a given gene with more than one probe set was represented by the average values of all its probe sets. Un-annotated probe sets were removed in the data pre-processing step. Only protein coding genes were considered in the analysis, as non-coding genes are often not well measured in microarray platforms. For RNAseq datasets, CPM (Count Per Million) were calculated to normalize the gene expression across samples and log_2_(CPM) were used for further analysis. Only protein coding genes were considered in the analysis to match the data in microarray datasets.

Transcriptome data were standardized by quantile-transformation to fit a normal distribution to avoid model misspecification when performing gene-level statistics. The expression values of all genes were normalized to the range of 0 to 1. Samples and genes with more than 30% missing values were removed from the analysis, and the remaining missing data were imputed using nearest neighbor averaging by the *impute.knn* function in the “impute” R package.

For all the datasets, covariates were manually annotated and curated based on the metadata available from the respective data sources. Datasets containing data from different tissues were separated into single tissues. To account for confounding sources of expression variations, the effects of known covariates, including age, gender, genotype, platform, disease, treatment, batch, etc, as well as hidden determinants of gene expression were estimated and removed by using PEER (probabilistic estimation of expression residuals) (Stegle et al. 2012), and the expression residuals were used for further analysis.

### Gene-Module Association Determination (G-MAD)

G-MAD makes use of the expression residuals of transcriptome datasets from large cohorts (datasets with over 80 samples). The expression levels of the gene-of-interest (target gene *T*) are used as a continuous trait to test whether a module M is enriched when *T* is highly expressed or, alternatively, whether it is depleted. The analysis uses the competitive gene set testing method CAMERA, which adjusts for inter-gene correlations (Wu and Smyth 2012). This adjustment is important, because left unadjusted too many significant results would emerge. To perform CAMERA, we first regress all genes *G* on *T* according to the following relationship

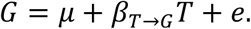

The fitting of this model equation to the observations is done separately for each data set by using the least squares method. The result is one fitted values *β*_*T*→*G*_ per gene. These coefficients define a set of statistics numerically characterizing the connection between the target gene *T* and any gene *G*. CAMERA provides a test of the null hypothesis that the average values of the *β* coefficients for the genes *G* in the module M are equal to the values for the genes not in the module. In order to correct for the inter-gene correlations a variance inflation factor is computed based on the average correlation coefficient 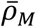 computed from the expression residuals obtained and only using the genes in the module M. When the average association scores between genes in the set and genes outside the set, 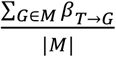 and 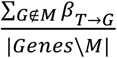, are compared on the final step, 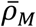 is included in the variance inflation factor. The resulting statistic revealing the association between the target gene *T* and M we refer to as the enrichment score *ES*_*M*_ (*T*).

The same procedure was conducted for all the genes in the analyzed datasets to obtain the enrichment *p*-value matrix between genes and modules in all the datasets. Two types of analyses can be applied on the gene-module *p*-value matrix. One can extract the *p*-values for one gene against all modules across the datasets to obtain the association between this gene and all modules; or extract the *p*-values for one module against all genes to check the association between this module and all genes. To avoid the situation where the final association scores are highly influenced by a few datasets with extremely low *p*-values, we converted the *p*-values to discrete association scores based on a significance threshold for each dataset. For the Bonferroni multiplicity correction, the significance thresholds for the *p*-values are either assessing genes for fixed modules 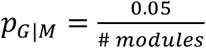 or assessing modules for fixed genes 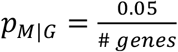. Gene-module associations with *p*-values that survived multiple testing corrections were set equal to 1 or −1, based on the enrichment direction, and 0 otherwise: 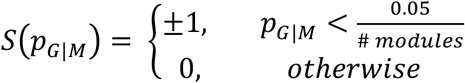, where *p*_*G*|*M*_ are one-sided *p*-values, corresponding to either positive or negative associations. The resulting *S*(*p*_*G*|*M*_) values were then meta-analyzed across the datasets, and the gene-module association scores (GMAS) were computed as the weighted averages of the scores with the weights functions of the sample sizes combined with the inter-gene correlation coefficients within modules. Denote *D*_*j*_, *j* = 1, …, *J* available datasets with corresponding sample sizes *n*_*j*_, *j* = 1, …, *J*, and average inter-gene correlations 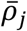, *j* = 1, …,*J*. Let the *p*-value obtained for the *j*^*th*^ dataset is *p*_*G*|*M*_(*j*). The final association score is then computed as

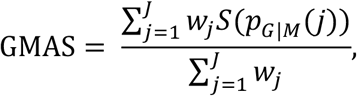

where weight for the *j*^*th*^ dataset is 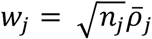. Under the null hypothesis, if we consider the positive and negative associations separately, the random variables *S*(*p*_*G*|*M*_(*j*)) follow a Bernoulli distribution with probability of success 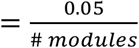. Therefore, statistic GMAS is the weighted sum of Bernoulli variables, whose theoretical distribution is hard to establish. The weight is proportional to the square root of the sample size in the *j*^*th*^ dataset. Another important component of *w*_*th*_ is the average correlation coefficients among genes in the module in the *j*^*th*^ dataset, 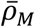, which reflects the co-expression or “level of activation” of the module for this dataset.

For the final decision we use thresholding of GMAS. We selected a very stringent threshold for GMAS, so that only a small proportion of the known gene-module connections are recovered. We found that a threshold of 0.268 enables us to recover 10% of the known gene-module links.

### Module-Module Association Determination (M-MAD)

M-MAD takes the association *p*-value matrix between a target module and all genes in all datasets (Fig. 2A bottom-left), and uses the −log_10_(*p*) values as a continuous trait to test whether other biological modules are enriched by containing genes that are highly associated with the target module. The analysis again uses the competitive gene set testing method CAMERA. Our function, −log10(*p*), transforms the *p*-values near zero to high positive values and *p*-values near 1 to transformed values near zero. Applied to *p*-values uniformly distributed in the interval between 0 and 1, the resulting transformed values have an exponential distribution skewed towards 0. CAMERA will then compute a *p*-value for testing the equality of the average transformed values for the genes in the other biological modules compared to all other genes. It will result in a small *p*-value when many of the genes in the other biological modules are relatively highly connected to the target module. The same analysis is performed for all modules to achieve a final association *p*-value matrix between modules. The Bonferroni correction was used to correct for the multiple testing errors with 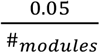 as the significance threshold. To avoid the situation where the final association scores are highly influenced by a few datasets with extreme low *p*-values, module-module connections with enrichment *p*-values that survived multiple testing corrections were allocated 1 or −1, based on the enrichment directions, and 0 otherwise. The results were then meta-analyzed across the datasets, and the module-module association scores (MMAS) were computed as the weighted averages of the connection scores by the sample sizes and inter-gene correlation coefficients within modules across datasets.

### Module network analysis

Module networks were constructed using Gephi 0.9.2 (Mathieu et al. 2009) based on either the module similarities or module connections from M-MAD. The Fruchterman-Reingold algorithm (Fruchterman and Reingold 1991) was used to create the network layout with a gravity value of 10. Iterations were stopped when the network reached stability. The node colors were obtained using the community detection algorithm (Vincent et al. 2008) embedded as the modularity tool in Gephi. Clusters with more than 20 nodes were colored to illustrate the module communities. The most frequent 10 biological terms (excluding biological meaningless words, such as “of”, “in”, or “and”) were used to represent the modules of these communities. The statistical characteristics of the module networks were computed using Gephi. For the network visualization of G-MAD results for one gene, modules were plotted according to their x and y coordinates of the module similarity network, and the gene-module association scores (GMAS) against all modules were used to color the modules using indicated color codes.

### Gene correlation network analysis

Gene correlation networks were constructed based on the Pearson correlation among genes of indicated modules in respective datasets using the “*layout_with_fr*” function in the *igraph* R package. Edges with correlation *p*-values lower than the indicated cutoffs in the figure panels were plotted.

### Cross validation

In order to test the predictive performance of G-MAD and compare it with the available methods using co-expression, we performed a cross validation analysis by removing groups of genes from modules, re-computing the associations between the removed genes and the reduced module and testing if we can rediscover the removed genes (Szklarczyk et al. 2016). We applied leave-one-out cross validation for modules with no more than 50 genes, and 10-fold cross validation for larger modules. The area under the receiver operating characteristic (ROC) curve (AUC) is used to estimate the performance of prediction, with an AUC of 1 indicating perfect prediction and 0.5 indicating random guess.

### Gene set enrichment analysis

Transcriptome data of uterus-specific *Arid1a* knockout mice(Kim et al. 2015) were downloaded from GEO under the accession number GSE72200. For enrichment analysis, genes were ranked based on their fold changes between *Arid1a* knockout and control samples, and gene set enrichment analysis (GSEA) was performed to identify the enriched gene sets using the R/fgsea package (Subramanian et al. 2005; Sergushichev 2016).

### Transcript-phenotype correlation analysis in mouse cohorts

Phenotype data, as well as transcriptome data of liver and white adipose tissue, from the BXD (Wu et al. 2014) and CTB6F2 (Schadt et al. 2008) mouse cohorts were downloaded from GeneNetwork (www.genenetwork.org). Spearman’s correlation coefficient *rho* was used to calculate the correlation between the transcript levels of ribosomal protein genes and metabolic phenotypes.

### Cell culture and siRNA transfection

Human embryonic kidney (HEK) 293 cells were cultured in DMEM supplemented with 10% fetal bovine serum, 100 IU/ml penicillin and 100 µg/ml streptomycin. HEK 293 cells were grown to approximately 70% confluence in 12-well plate. The cells were treated with either scrambled siRNA, or human *DDT* / *BOLA3* siRNA (Dharmacon) mixed with lipofectamine 2000 to yield a final concentration of 100nM according to the supplier’s protocol. After siRNA treatment for 48 hours, cells were collected for quantitative real-time PCR assay. Primers used in this assay are listed in Supplemental Table S5. Statistical significance was determined by two-tailed Student’s t-test.

### Mitochondrial function assay

Mitochondrial oxygen consumption rate (OCR) was measured on a Seahorse XFe96 analyzer (Agilent) according to the manufacturer’s protocol. HEK 293 cells were seeded on to 96-well XF analyzer assay plate. Cells were treated with scrambled siRNA or human *DDT / BOLA3* siRNA. After 48 hours siRNA treatment, Seahorse XFe96 analyzer was used to measure OCR of the cells. After basal OCR levels were measured, HEK 293 cells were cumulatively treated with 1µM Oligomycin (ATP synthase inhibitor), then 3µM carbonyl cyanide 4-(trifluoromethoxy) phenylhydrazone (FCCP, mitochondrial uncoupler). Then, a mixture of 1µM Antimycin A (mitochondrial respiratory chain Complex III inhibitor) and 1µM Rotenone (Complex I inhibitor) was added. OCR levels were normalized to total protein content per well determined by Lowry protein assay. Statistical significance was determined by two-tailed Student’s t-test.

### *C. elegans* experiments

Lipid droplets were stained in *C. elegans* as described previously (Li et al. 2018). Inhibition of ribosome in early stage of worms affects their development and growth, so RNAi was performed after the worms reached adulthood. Specifically, L1 larvae of N2 worms were grown on regular nematode growth media (NGM) plates at 20°C for 2 days until reaching adulthood. Then worms were then transferred to RNAi plates with 1mM IPTG containing HT115 bacteria expressing RNAi clones for ribosomal genes or empty vector. After 2 days of RNAi treatment, worms were collected, washed twice with 1x PBS and then suspended in 120 µl of PBS. Then 120 µl 2x MRWB buffer (160 mM KCl, 40 mM NaCl, 14 mM Na_2_EGTA, 30 mM PIPES pH 7.4, 1 mM Spermidine, 0.4 mM Spermine, 2% paraformaldehyde, 0.2% beta-mercaptoethanol) was added. The worms were taken through 3 freeze-thaw cycles between dry ice/ethanol mixture and warm running tap water, followed by 1 minute spinning at 14,000g. Worms were then washed once using PBS to remove paraformaldehyde. Oil Red O staining of lipid droplets was performed after fixation. Worms were re-suspended and dehydrated in 60% isopropanol. 250 µl of 60% Oil Red O stain was added to each sample, and samples were incubated overnight at room temperature. Worms were washed twice in 60% isopropanol solution after Oil Red O staining. The region immediately behind the pharynx of each worm was used for imaging of the lipid droplets (Li et al. 2018). The lipid droplets were quantified using Fiji (ImageJ) as previously described (Li et al. 2018). Statistical significance was determined by two-tailed Student’s t-test.

## Data access

### Data Availability

All data included in the study is available from https://systems-genetics.org/.

### Code Availability

Code used in the study is available from https://github.com/lihaone/GeneBridge.

## Acknowledgments

We are grateful to the research groups who made these data publicly available for systems biology research. We thank N. Agarwal for help in data preprocessing. We thank the entire J.A. lab for comments and discussions. H.L. is the recipient of a doctoral scholarship from the China Scholarship Council. This work was supported by grants from the EPFL, the ERC (AdG-787702), the SNSF (310030B-160318), the AgingX program of the Swiss Initiative for Systems Biology (RTD 2013/153), and the NIH (R01AG043930).

## Author contributions

Conceptualization: H.L., and J.A.; Data curation: H.L.; Formal analysis: H.L., D.R., S.M., and J.A.; Funding acquisition: R.W.W., M.R-R., K.S., and J.A.; Methodology: H.L., D.R., A.K., Q.H., S.M., and J.A.; Resources: H.L., D.R., F.P.A.D., M.B.S., S.M., and J.A.; Software: H.L., and F.P.A.D.; Validation: H.L., T.Y.L., C-M.O., A.W.G., and E.K.; Visualization: H.L., and J.A.; Writing – original draft: H.L., and J.A.; Writing – review & editing: H.L., S.M., and J.A.

---

**Fig. S1.**
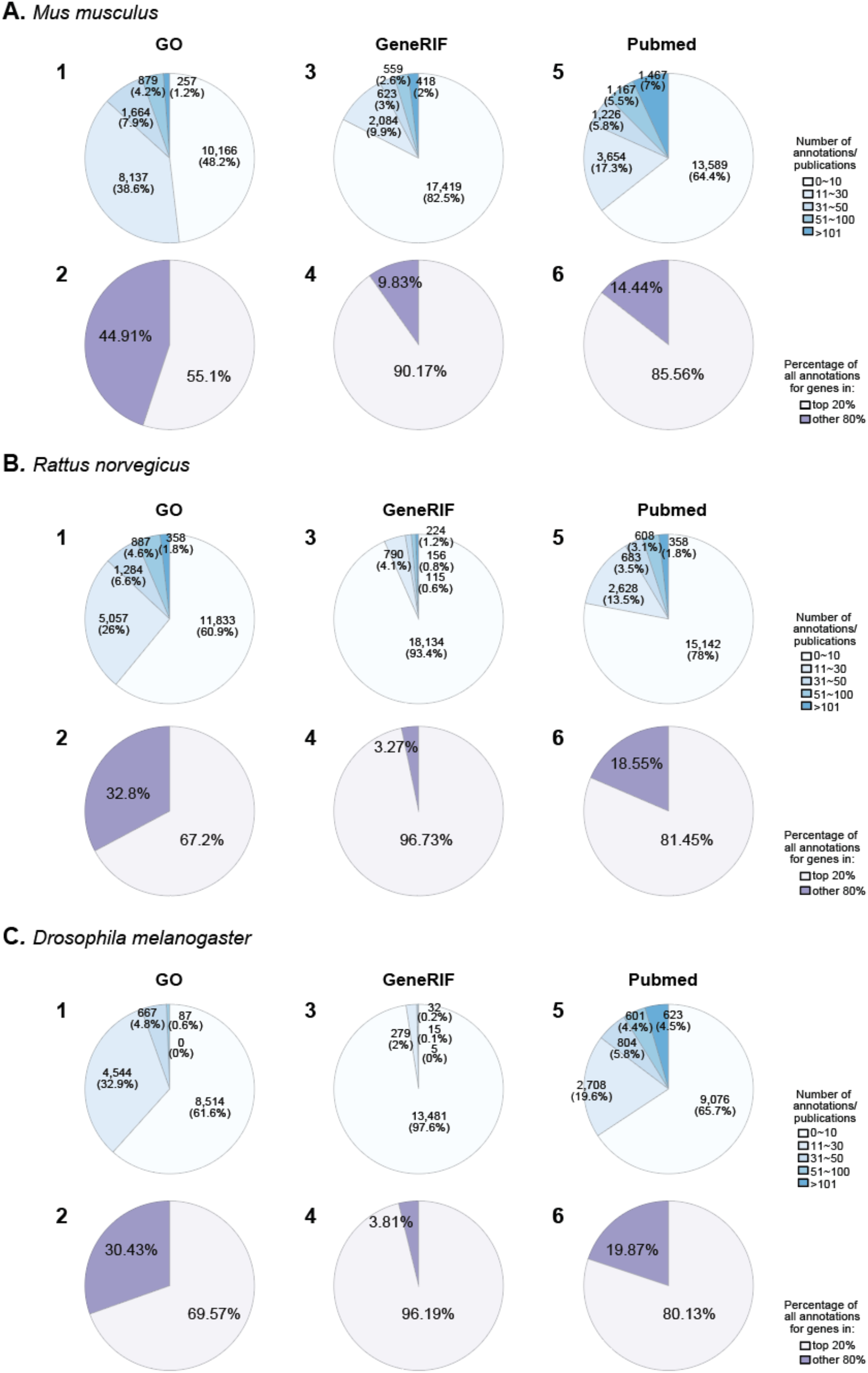
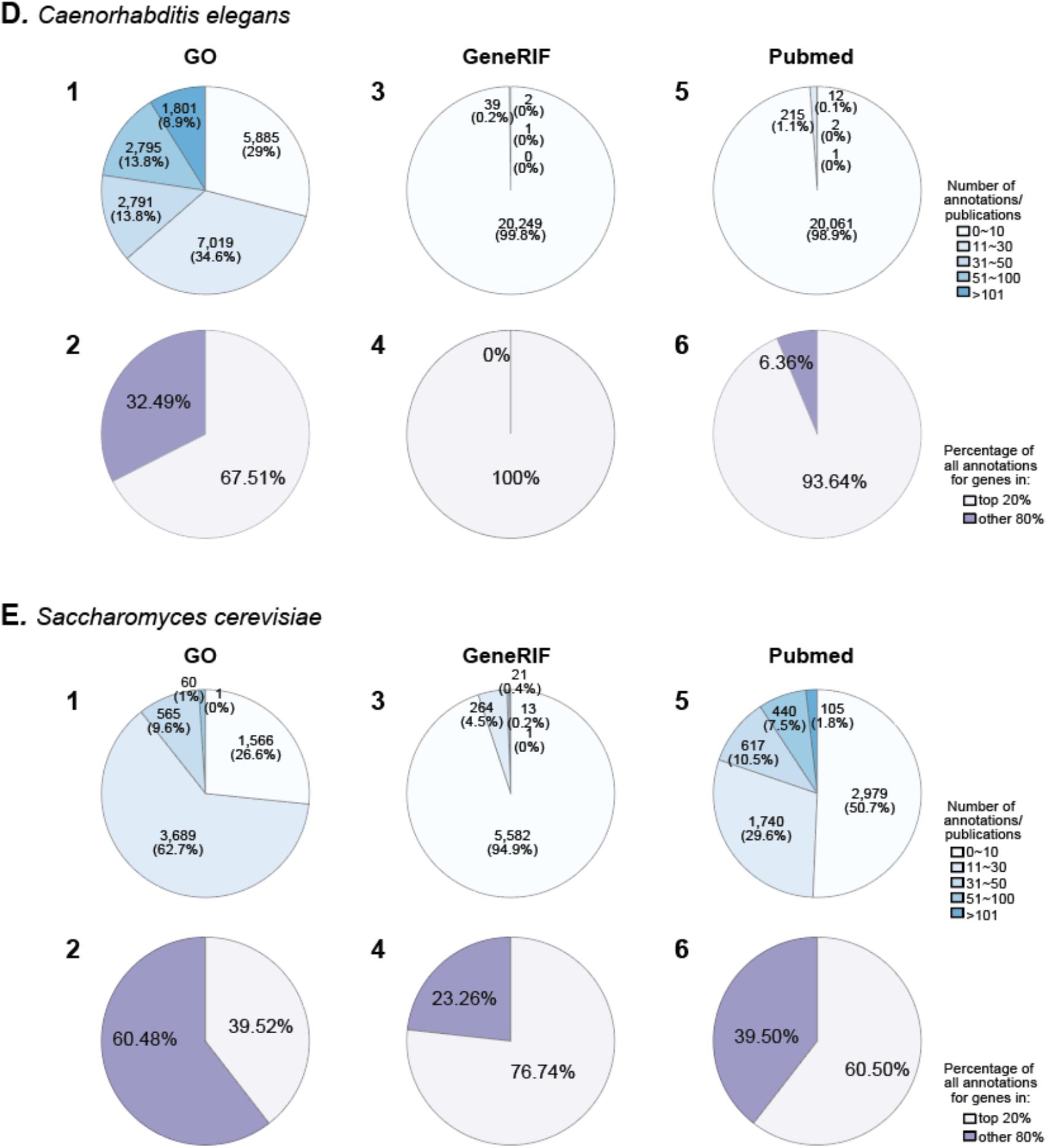
Statistical summary of available annotations for genes in *M. musculus* (A), *R. norvegicus* (B), *D. melanogaster* (C), *C. elegans* (D), and *S. cerevisiae* (E). The number of annotations per gene in GO (**1**), GeneRIF (**3**), and the number of publications in PubMed (**5**) for respective species. The percentage of the annotations/ publications covering the top 20% most annotated genes (**2**, **4**, **6**).

**Fig. S2.**
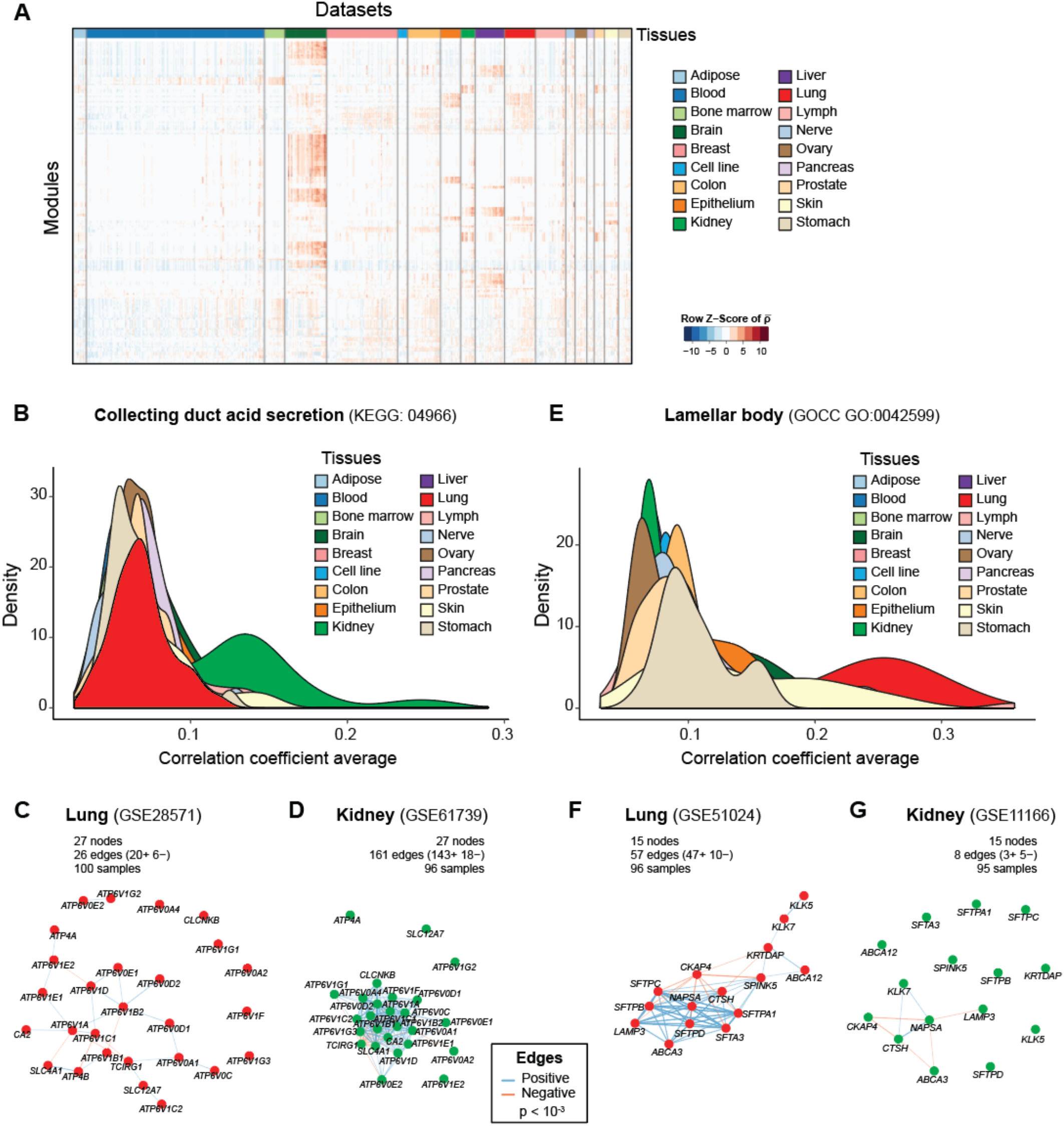
Tissue specific co-expression of modules. **A**, Heatmap showing the correlation coefficient averages of genes 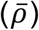 in modules from expression data of a subset of all human datasets. Datasets from different tissues are arranged and colored (top bar). Modules are clustered in rows using hierarchical clustering. 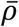 values for each module are centered and scaled per module. **B**, **E**, Distribution of the co-expression of genes in the “collecting duct acid secretion” (**B**) or “lamellar body” (**E**) module across tissues in human. The average correlation coefficient of the gene pairs of the module in 1,300 human expression datasets from 18 major tissues was used as to illustrate the co-expressions of this module across tissues. Genes in “collecting duct acid secretion” (**B**) and “lamellar body” (**E**) module have higher co-expression in datasets from kidney and lung, respectively, indicating the potential to assign tissue-specificity. **C**-**G**, Pearson correlation network of genes in the “collecting duct acid secretion” (**C**, **D**) or “lamellar body” (**F**, **G**) module in representative datasets of lung (**C**, **F**) and kidney (**D**, **G**). The number of genes (nodes) and gene pairs (edges) that survive the indicated threshold of correlation significance are shown. Genes in the “collecting duct acid secretion” module have higher co-expression in datasets from the kidney (**C**) than the lung (**D**), while genes in the “lamellar body” module have higher co-expression in datasets from the lung (**F**) than the kidney (**G**).

**Fig. S3.**
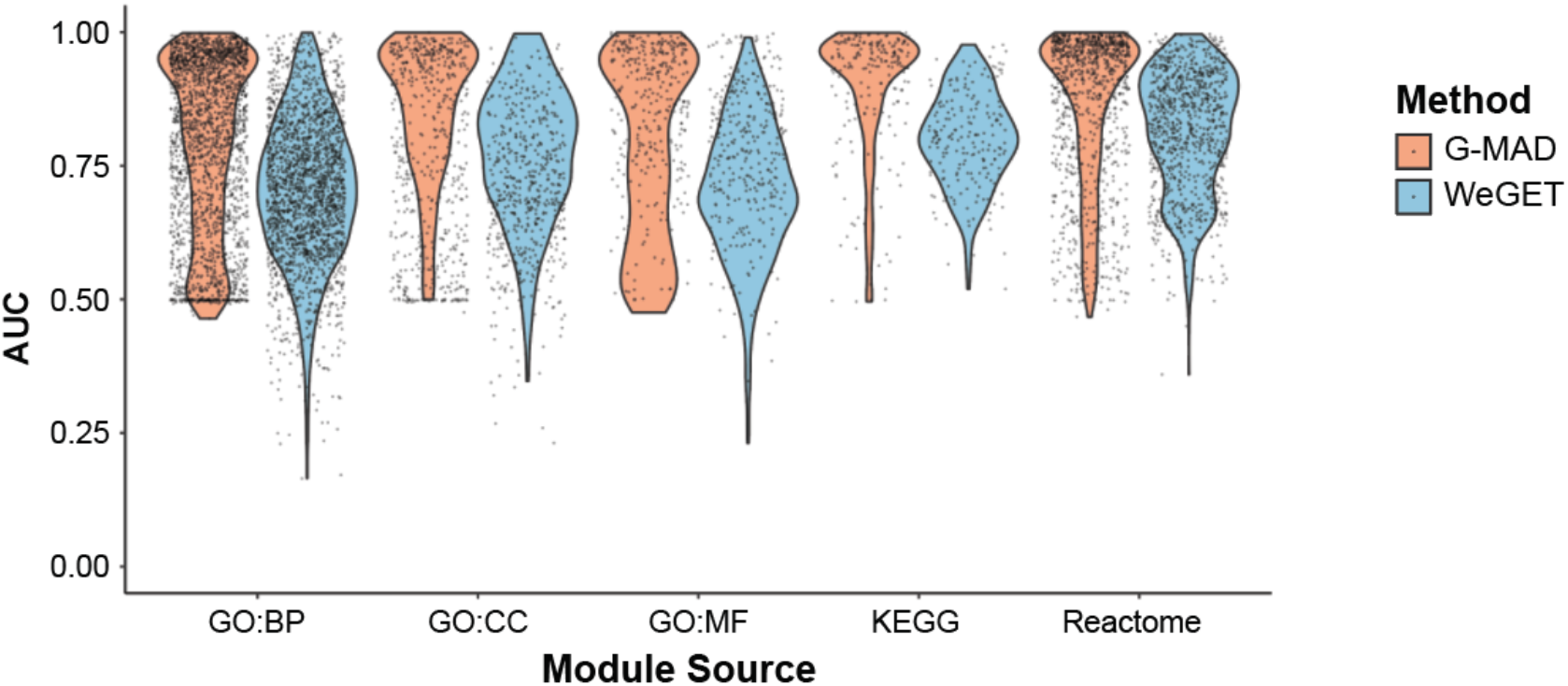
Comparison of the predictive performance of G-MAD with available methods. The predictive performance of G-MAD is compared to WeGET using cross-validation. Cross validation evaluates the robustness of the methods by removing the genes from the query module and test the performance in redetecting them. Performance of the method is computed as the area under the receiver operating characteristic curve (AUC) for each module. A high AUC indicates that most of the genes in the module are rediscovered when they are removed from the module in the analysis.

**Fig. S4.**
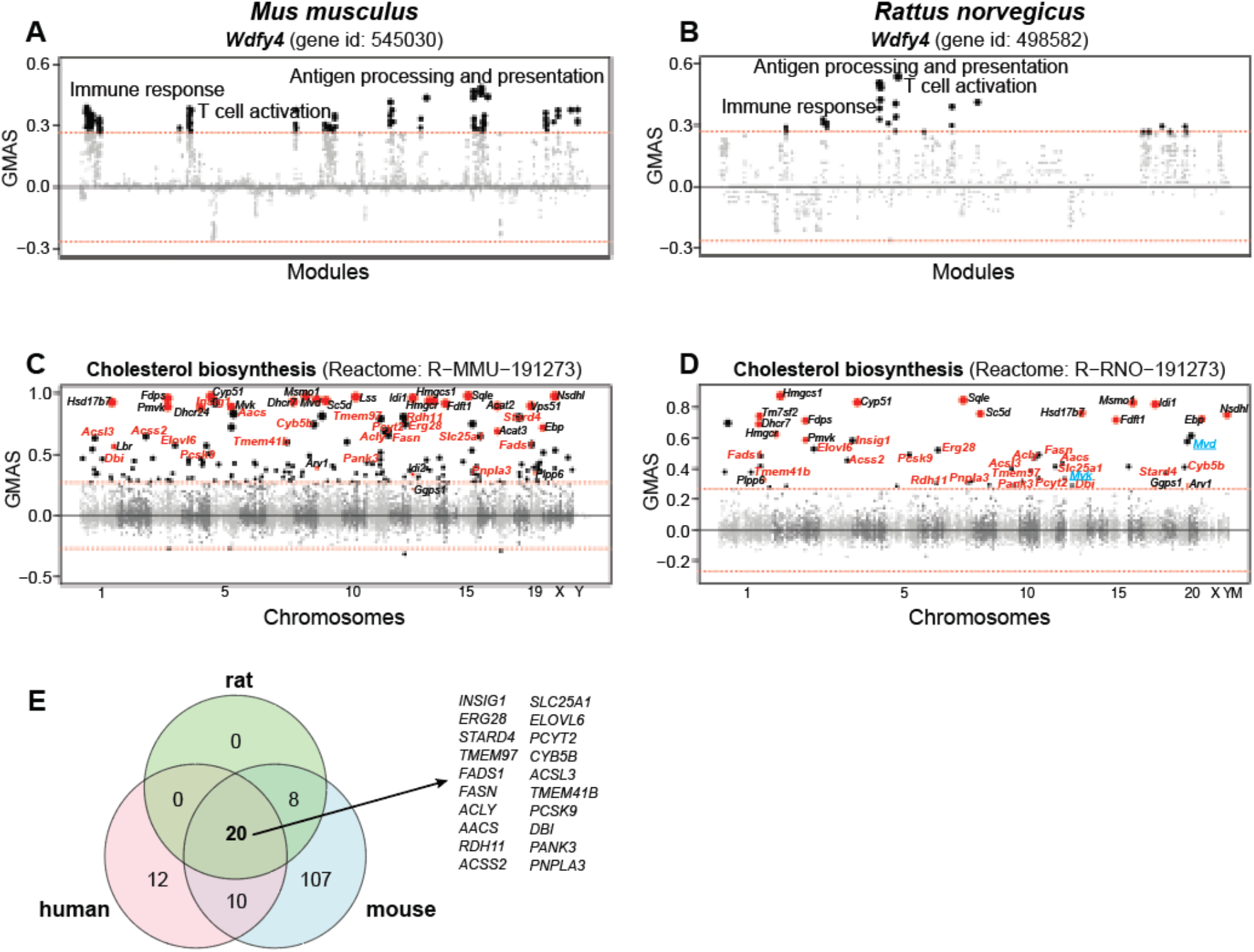
G-MAD in mouse and rat confirms the gene-module connections between *WDFY4* and T cell activation, and links 20 new genes with cholesterol biosynthesis. **A, B**, G-MAD of *Wdfy4* in mouse (**A**) and rat (**B**) confirms its involvement in T cell activation and immune response. The threshold of significant gene-module association is indicated by the red dashed line. Modules are organized by the module similarities. Known modules connected to *Wdfy4* from annotations are shown in red dots (no connected modules for *Wdfy4*), and other modules with GMAS over the threshold are shown with black dots. **C, D**, G-MAD confirms the involvement of novel genes in cholesterol biosynthesis in mouse (**C**) and rat (**D**). The threshold of significant gene-module association is indicated by the red dashed line. Genes are arranged based on their genetic positions. Genes annotated to be involved in cholesterol biosynthesis are shown in red dots, and genes with GMAS over 0.268 are shown in black dots. Novel genes conserved in human, mouse and rat are highlighted in red bold text. *Mvd* and *Mvk* (highlighted in blue text in **D**) are included in the annotation of cholesterol biosynthesis module in human and mouse, but not in rat. **E**, Venn diagram comparing G-MAD results of cholesterol biosynthesis in human, mouse, and rat. 20 novel genes were identified with conserved associations with cholesterol biosynthesis in all 3 species.

**Fig. S5.**
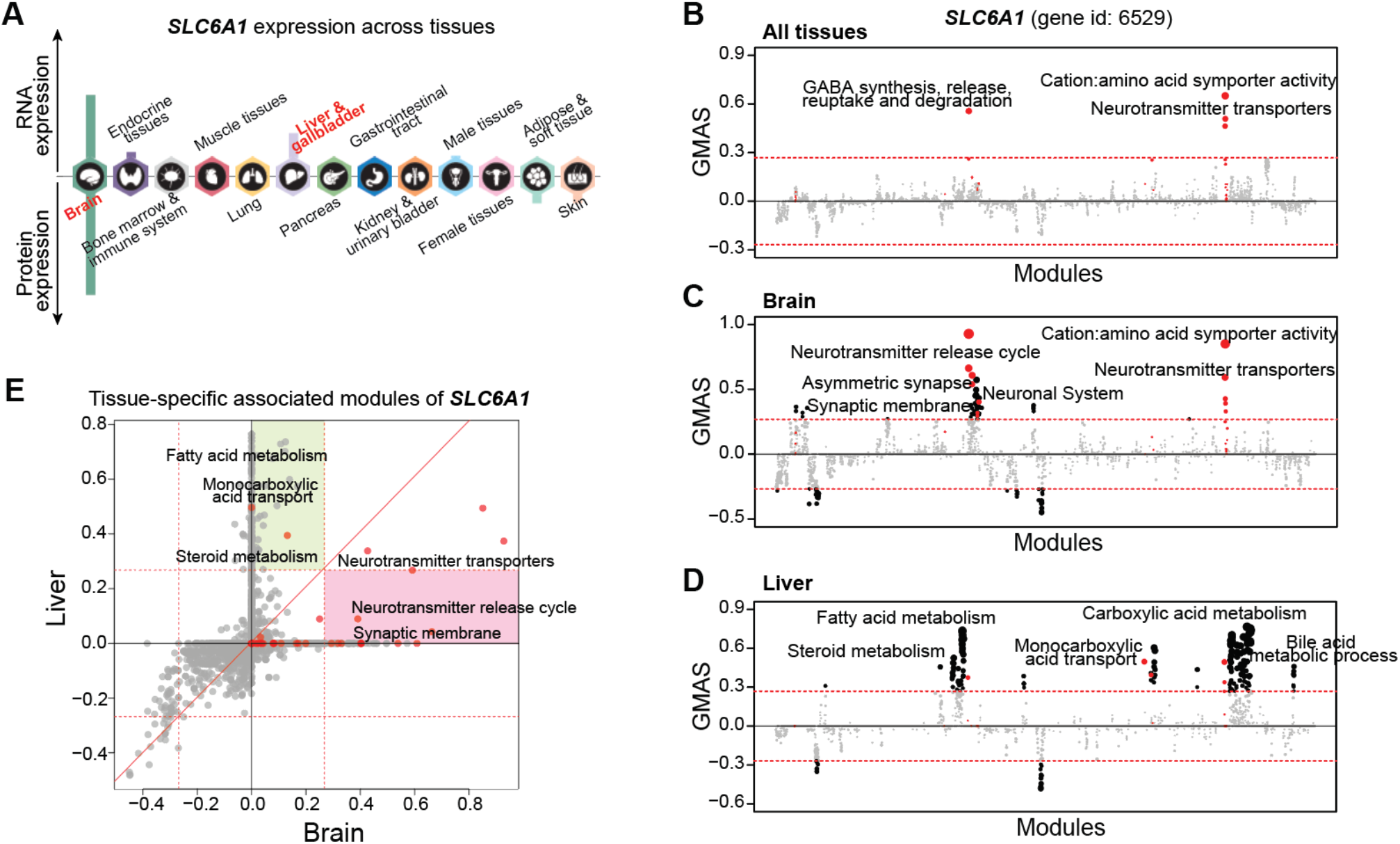
G-MAD identifies tissue-specific associated modules for *SLC6A1* by using datasets from different tissues. **A**, Expression patterns of *SLC6A1* across tissues. The figure was adapted from the Human Protein Atlas. **B**-**D**, G-MAD of *SLC6A1* in human using datasets from all tissues (**B**), from brain (**C**), or from liver (**D**). The threshold of significant gene-module association is indicated by the red dashed line. Modules are organized by their similarities. Known modules connected to *SLC6A1* from gene annotations are shown in red dots, and other modules with GMAS over the threshold are shown by black dots. **E**, Comparison of G-MAD results of *SLC6A1* in brain and liver. Known modules connected to *SLC6A1* are shown in red dots. The threshold of significant gene-module association is indicated by the red dashed line. Modules significantly associated with *SLC6A1* only in one specific tissue are highlighted. The comparison of the association results of *SLC6A1* in brain and liver can be found at Supplemental Table S4.

**Fig. S6.**
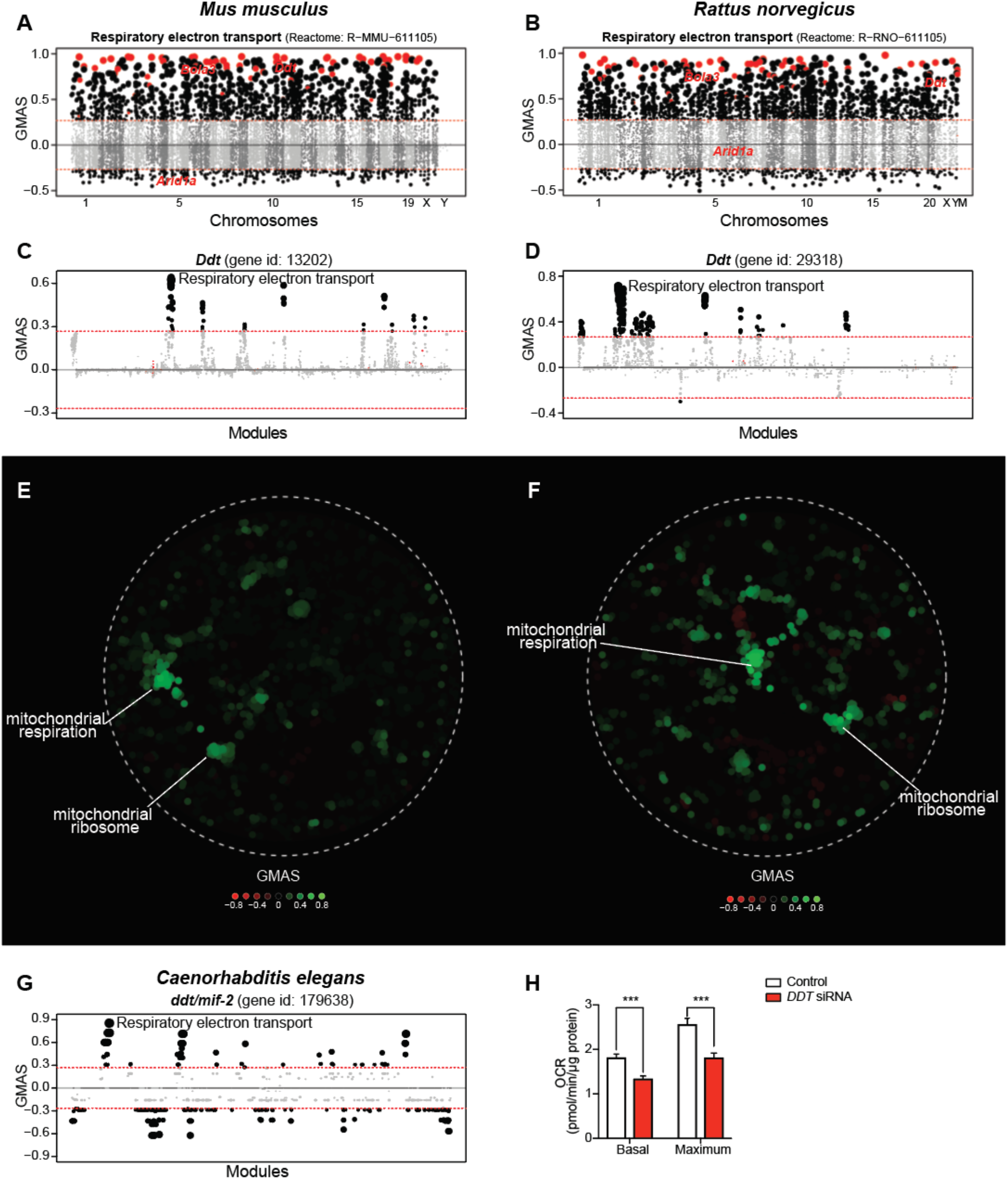
G-MAD verifies the potential involvement of *DDT* in mitochondrial respiration in mouse and rat. **A, B**, G-MAD of respiratory electron transport in mouse (**A**) and rat (**B**). The threshold of significant gene-module association is indicated by the red dashed line. Genes are arranged based on their genetic positions. Genes annotated to be involved in respiratory electron transport are shown in red dots, and other genes with GMAS over 0.268 are highlighted by black dots. **C, D**, G-MAD of *Ddt* in mouse (**C**) and rat (**D**) confirms its involvement in mitochondrial respiratory chain. The threshold of significant gene-module association is indicated by the red dashed line. Modules are organized by their similarities. Known modules connected to *Ddt* from gene annotations are shown in red dots, and other modules with GMAS over the threshold are shown by black dots. **E, F**, Network plots showing the significantly connected modules of *Ddt* in mouse (**E**) and rat (**F**). Modules are colored based on their GMAS against *Ddt*. **G**, G-MAD of *ddt/mif-2* in *C. elegans* confirms its involvement in mitochondrial respiratory chain function also in invertebrates. The threshold of significant gene-module association is indicated by the red dashed line. Modules are organized by their similarities. Known modules connected to *ddt/mif-2* from gene annotations are shown in red dots, and other modules with GMAS over the threshold are shown by black dots. **H**, *DDT* RNAi reduced basal and maximum oxygen consumption rate (OCR) in HEK293 cells. Results were computed from Fig. **4F**. ***, *p* < 0.001.

**Fig. S7.**
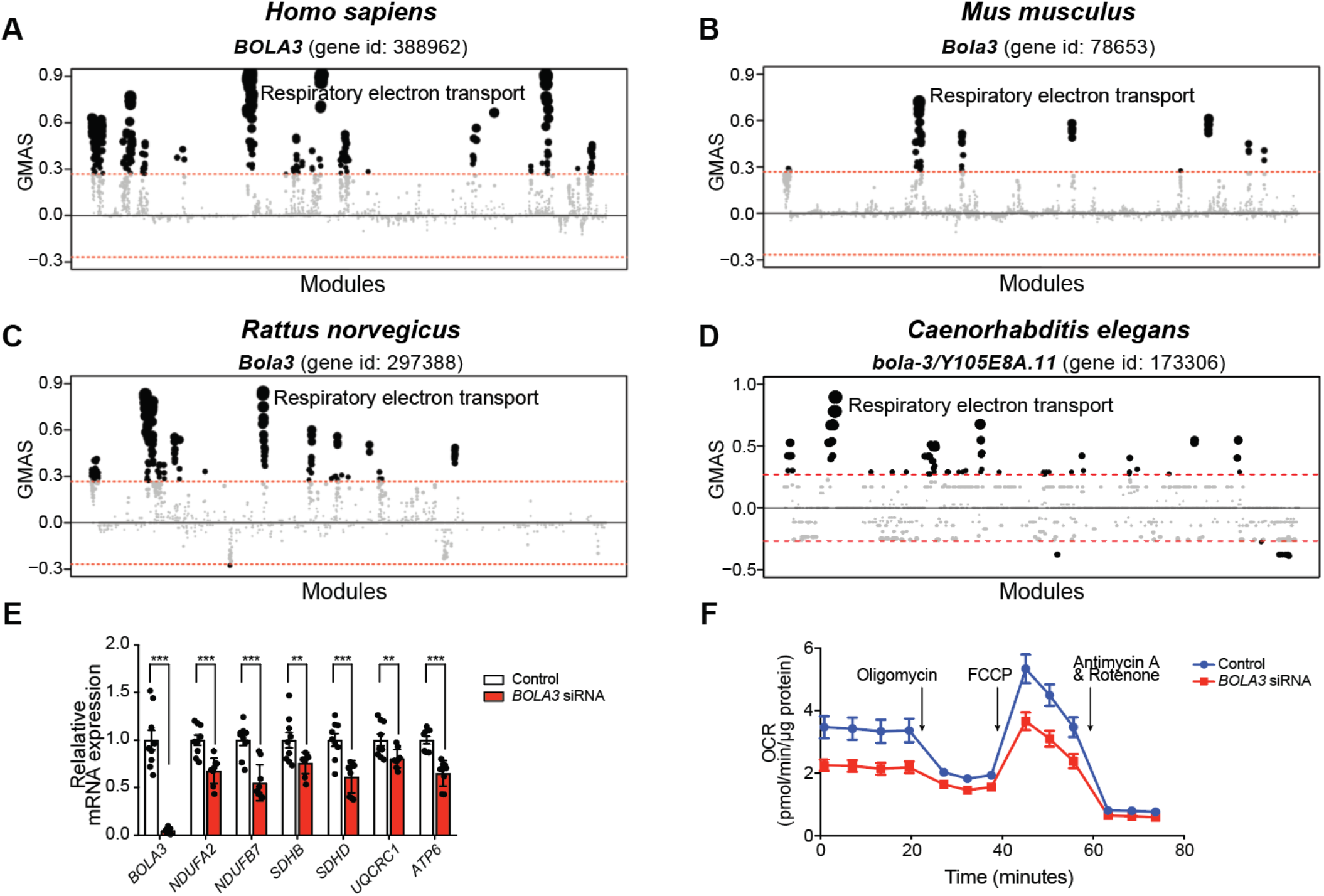
G-MAD confirms the involvement of *BOLA3* in mitochondrial respiration. **A-D**, *BOLA3/Bola3/bola-3* associates with mitochondrial respiratory chain modules in human (**A**), mouse (**B**), rat (**C**), and *C. elegans* (**D**). The threshold of significant gene-module association is indicated by the red dashed line. Modules are organized by module similarities. Known modules connected to *BOLA3/Bola3/bola-3* from annotations are highlighted in red (no connected modules for *BOLA3/Bola3/bola-3*), and other modules with GMAS over the threshold are colored in black. Dot sizes reflect the GMAS of *BOLA3* against the respective modules. **E**, Silencing *BOLA3* expression in HEK293 cells decreases expression levels of indicated genes involved in mitochondrial respiratory chain complexes. Error bars represent standard errors. *, *p* < 0.05; **, *p* < 0.01; ***, *p* < 0.001. n=9. **F**, *BOLA3* knockdown leads to the reduction of oxygen consumption rate (OCR) as a reflection of mitochondrial respiration in human HEK293 cells. Addition of specific mitochondrial inhibitors, including the oligomycin (ATPase inhibitor), FCCP (uncoupling agent), and rotenone/antimycin A (electron transport chain inhibitors) are indicated.

**Fig. S8.**
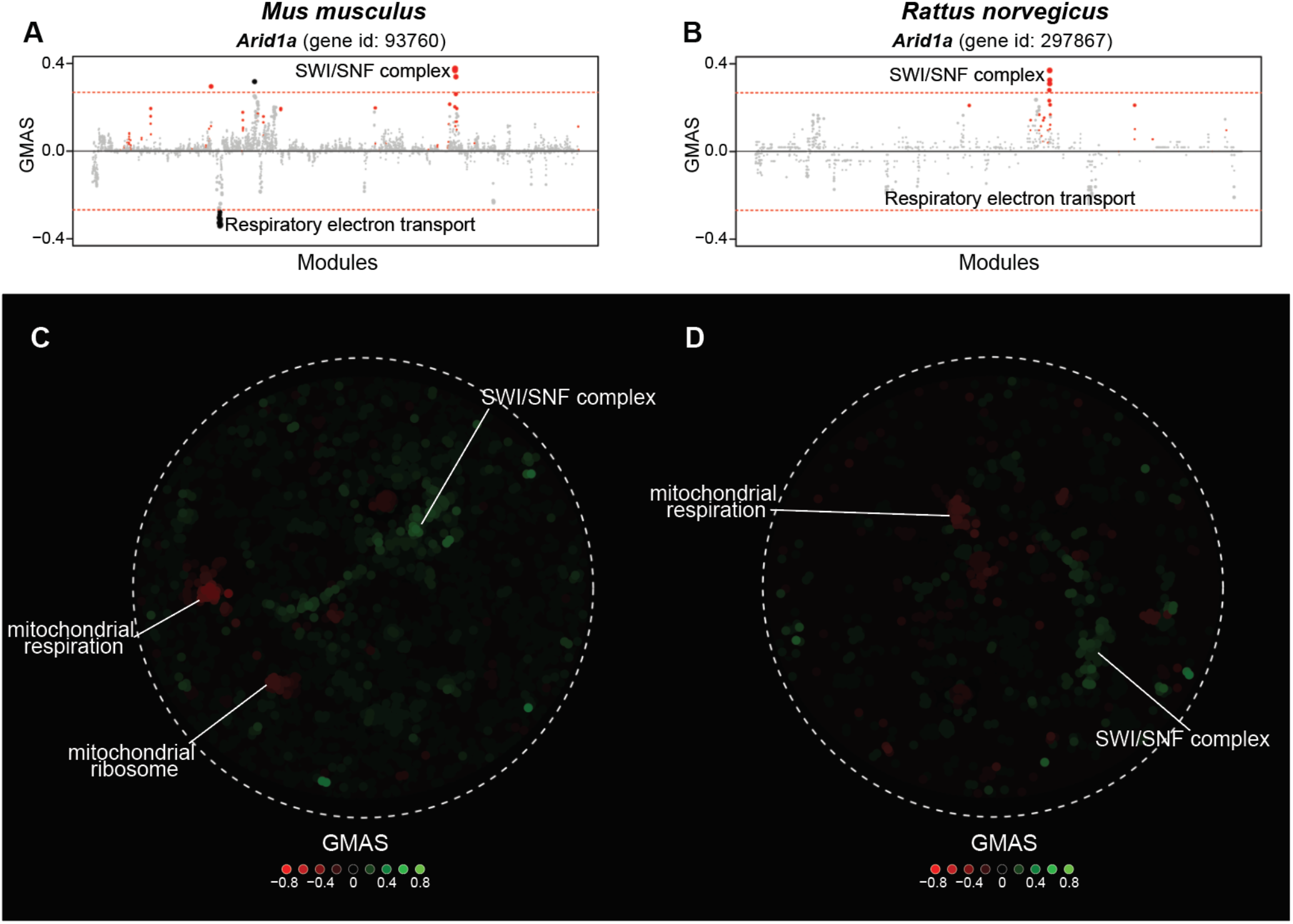
G-MAD verifies the negative association of *Arid1a* with mitochondrial respiration in mouse and rat. **A, B**, G-MAD of *Arid1a* in mouse (**A**) and rat (**B**) confirms its negative association with mitochondrial respiratory chain. The threshold of significant gene-module association is indicated by the red dashed line; note that the associations are only suggestive in the mouse. Modules are organized by their similarities. Known modules connected to *Arid1a* from gene annotations are shown in red dots, and other modules with GMAS over the threshold are shown by black dots. **C, D**, Network plot showing the significantly connected modules of *Arid1a* in mouse (**C**) and rat (**D**). Modules are colored based on their GMAS against *Arid1a*.

**Fig. S9.**
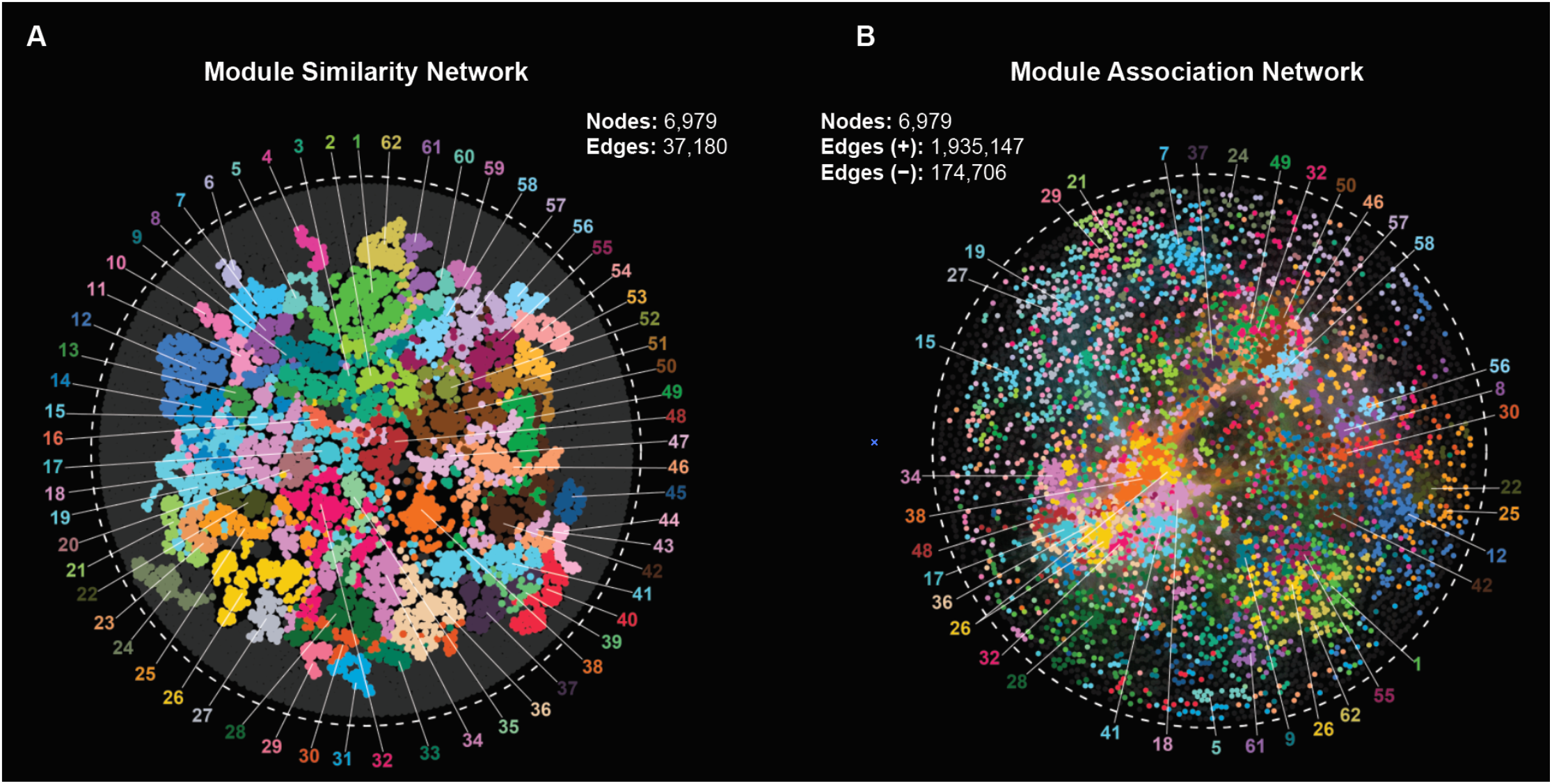
Comparison of module similarity network and module association network. Module network obtained from module similarity (**A**) and association (**B**; M-MAD) were put together to facilitate the comparison.

**Fig. S10.**
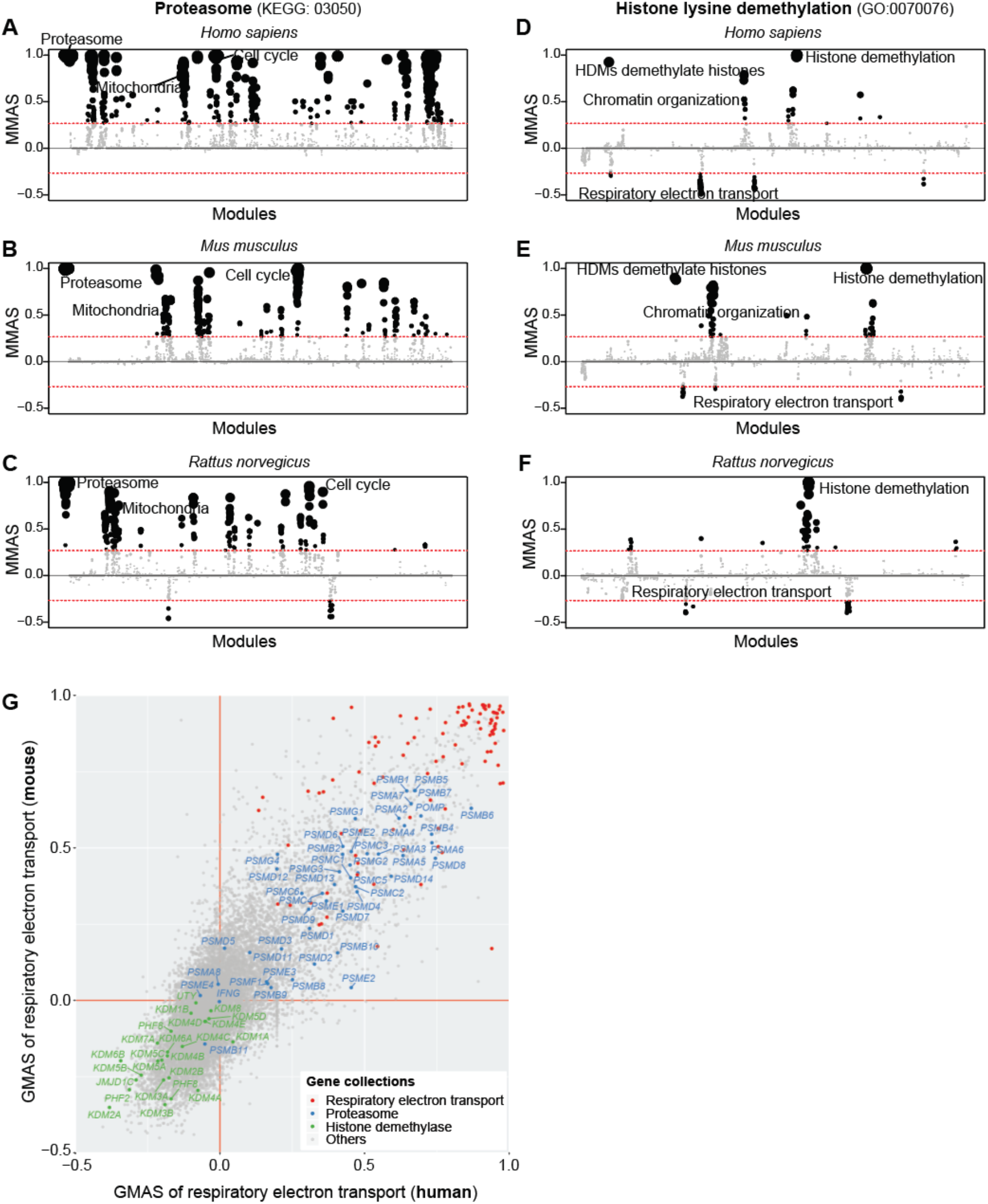
M-MAD and G-MAD identify the connections between respiratory electron transport chain and proteasome, as well as histone lysine demethylation. **A-F**, M-MAD results for proteasome (**A-C**) and histone lysine demethylation (**D-F**) in human (**A, D**), mouse (**B, E**), and rat (**C, F**). The threshold of significant module-module connection is indicated by the red dashed line. Modules are organized by the module similarities. Dot sizes are proportional to MMAS of the respective modules. **G**, G-MAD results for electron transport chain from human and mouse are plotted in the x and y axes, respectively. Genes annotated to be involved in respective modules are indicated in different colors.

**Fig. S11.**
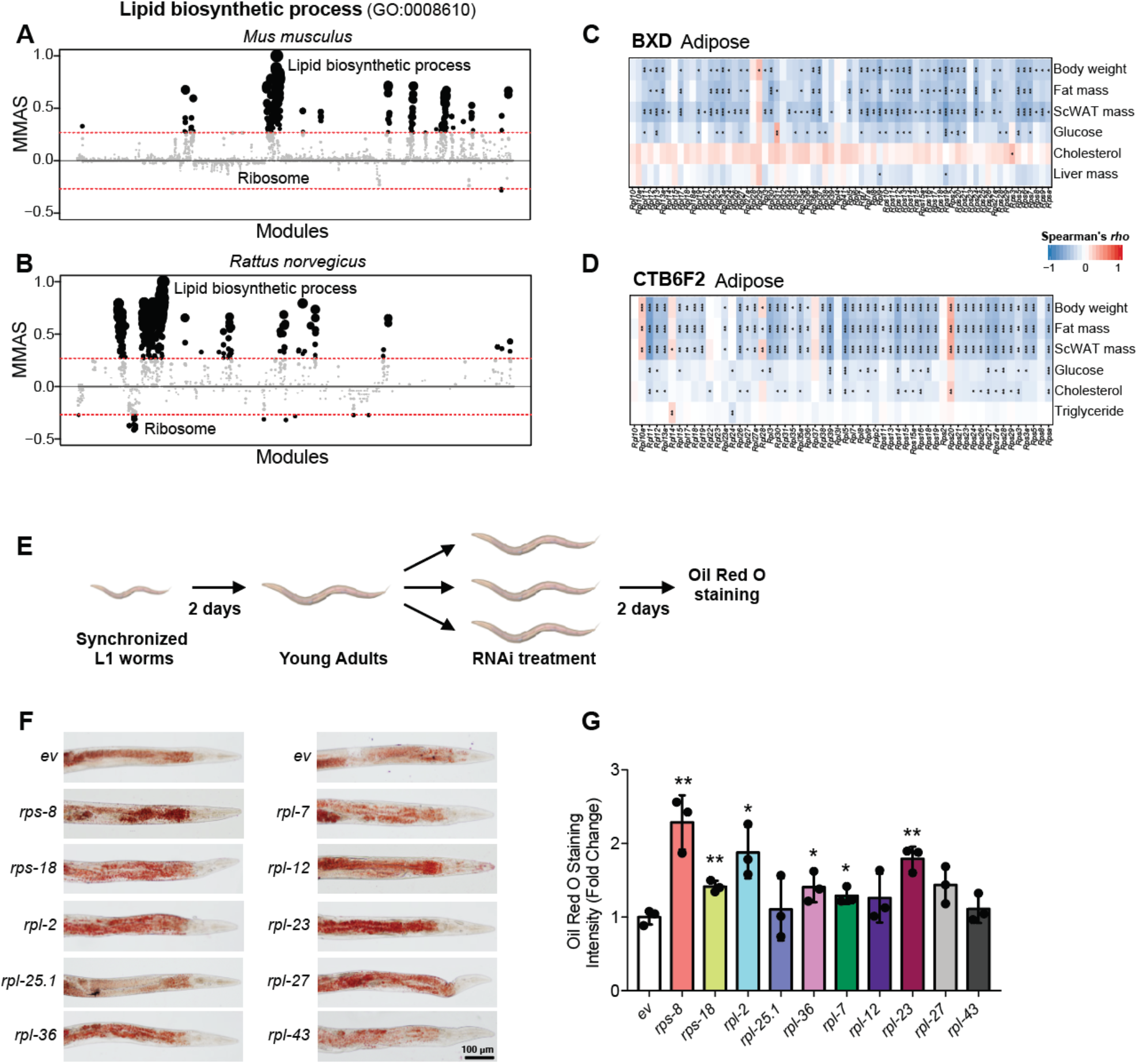
Validations of the negative association between ribosome and lipid biosynthetic modules. **A-B**, M-MAD results for lipid biosynthetic process in mouse (**A**) and rat (**B**). The threshold of significant module-module connection is indicated by the red dashed line; note that the associations are only suggestive in the mouse. Modules are organized by the module similarities. Dot sizes are proportional to MMAS of the respective modules. **C-D**, Transcripts of genes encoding for ribosomal proteins in the white adipose tissue negatively correlate with metabolic traits in the BXD (**C**) and CTB6F2 (**D**) mouse cohorts. *, *p* < 0.05; **, *p* < 0.01; ***, *p* < 0.001. ScWAT, subcutaneous white adipose tissue. **E**, Scheme of the experimental design. L1 worm larvae were grown on regular NGM plates at 20°C for 2 days and then transferred to RNAi plates with 1mM IPTG containing HT115 bacteria expressing RNAi clones for ribosomal genes or empty vector (*ev*). After 2 days, worms were collected and lipid droplets were stained using Oil Red O. **F**, Representative images of worm lipid droplet staining after RNAi of ribosomal genes. **G**, Quantification of the lipid droplet staining intensity in **F**. *, *p* < 0.05; **, *p* < 0.01; ***, *p* < 0.001. n=3.

**Supplemental Table S1.** Datasets used in this study.

**Supplemental Table S2.** Top 10 most frequent terms of the module clusters in Fig. 2C.

**Supplemental Table S3.** Tissue-specific association of *EHHADH* in kidney and liver tissues in humans.

**Supplemental Table S4.** Tissue-specific association of *SLC6A1* in brain and liver tissues in humans.

**Supplemental Table S5.** G-MAD results on respiratory electron transport in human, mouse and rat.

**Supplemental Table S6.** qPCR primer sequences used in this study.

## References

Arroyo JD, Jourdain AA, Calvo SE, Ballarano CA, Doench JG, Root DE, Mootha VK. 2016. A Genome-wide CRISPR Death Screen Identifies Genes Essential for Oxidative Phosphorylation. Cell metabolism 24(6): 875–885.

Ashburner M, Ball CA, Blake JA, Botstein D, Butler H, Cherry JM, Davis AP, Dolinski K, Dwight SS, Eppig JT et al. 2000. Gene ontology: tool for the unification of biology. The Gene Ontology Consortium. Nature genetics 25(1): 25–29.

Austin CP, Battey JF, Bradley A, Bucan M, Capecchi M, Collins FS, Dove WF, Duyk G, Dymecki S, Eppig JT et al. 2004. The knockout mouse project. Nature genetics 36(9): 921–924.

Barabasi AL, Gulbahce N, Loscalzo J. 2011. Network medicine: a network-based approach to human disease. Nature reviews Genetics 12(1): 56–68.

Barrett T, Wilhite SE, Ledoux P, Evangelista C, Kim IF, Tomashevsky M, Marshall KA, Phillippy KH, Sherman PM, Holko M et al. 2013. NCBI GEO: archive for functional genomics data sets--update. Nucleic acids research 41(Database issue): D991–995.

Bartz F, Kern L, Erz D, Zhu M, Gilbert D, Meinhof T, Wirkner U, Erfle H, Muckenthaler M, Pepperkok R et al. 2009. Identification of cholesterol-regulating genes by targeted RNAi screening. Cell metabolism 10(1): 63–75.

Bastian F, Parmentier G, Roux J, Moretti S, Laudet V, Robinson-Rechavi M. 2008. Bgee: Integrating and Comparing Heterogeneous Transcriptome Data Among Species. In Data Integration in the Life Sciences, (ed. A Bairoch, S Cohen-Boulakia, C Froidevaux), pp. 124–131. Springer Berlin Heidelberg, Berlin, Heidelberg.

Bogue MA, Grubb SC, Walton DO, Philip VM, Kolishovski G, Stearns T, Dunn MH, Skelly DA, Kadakkuzha B, TeHennepe G et al. 2018. Mouse Phenome Database: an integrative database and analysis suite for curated empirical phenotype data from laboratory mice. Nucleic acids research 46(D1): D843–D850.

Calvo SE, Clauser KR, Mootha VK. 2016. MitoCarta2.0: an updated inventory of mammalian mitochondrial proteins. Nucleic acids research 44(D1): D1251–1257.

Cameron JM, Janer A, Levandovskiy V, Mackay N, Rouault TA, Tong WH, Ogilvie I, Shoubridge EA, Robinson BH. 2011. Mutations in iron-sulfur cluster scaffold genes NFU1 and BOLA3 cause a fatal deficiency of multiple respiratory chain and 2-oxoacid dehydrogenase enzymes. American journal of human genetics 89(4): 486–495.

Carroll RG, Zaslona Z, Galvan-Pena S, Koppe EL, Sevin DC, Angiari S, Triantafilou M, Triantafilou K, Modis LK, O’Neill LA. 2018. An unexpected link between fatty acid synthase and cholesterol synthesis in proinflammatory macrophage activation. The Journal of biological chemistry 293(15): 5509–5521.

Carvill GL, McMahon JM, Schneider A, Zemel M, Myers CT, Saykally J, Nguyen J, Robbiano A, Zara F, Specchio N et al. 2015. Mutations in the GABA Transporter SLC6A1 Cause Epilepsy with Myoclonic-Atonic Seizures. American journal of human genetics 96(5): 808–815.

Chesler EJ, Lu L, Wang J, Williams RW, Manly KF. 2004. WebQTL: rapid exploratory analysis of gene expression and genetic networks for brain and behavior. Nature neuroscience 7(5): 485–486.

Costanzo M, Baryshnikova A, Bellay J, Kim Y, Spear ED, Sevier CS, Ding H, Koh JL, Toufighi K, Mostafavi S et al. 2010. The genetic landscape of a cell. Science 327(5964): 425–431.

Croft D, O’Kelly G, Wu G, Haw R, Gillespie M, Matthews L, Caudy M, Garapati P, Gopinath G, Jassal B et al. 2011. Reactome: a database of reactions, pathways and biological processes. Nucleic acids research 39(Database issue): D691–697.

D’Amico D, Sorrentino V, Auwerx J. 2017. Cytosolic Proteostasis Networks of the Mitochondrial Stress Response. Trends in biochemical sciences 42(9): 712–725.

Dickinson ME, Flenniken AM, Ji X, Teboul L, Wong MD, White JK, Meehan TF, Weninger WJ, Westerberg H, Adissu H et al. 2016. High-throughput discovery of novel developmental phenotypes. Nature 537(7621): 508–514.

Dolgin E. 2017. The most popular genes in the human genome. Nature 551(7681): 427–431.

Edwards AM, Isserlin R, Bader GD, Frye SV, Willson TM, Yu FH. 2011. Too many roads not taken. Nature 470(7333): 163–165.

Eisen MB, Spellman PT, Brown PO, Botstein D. 1998. Cluster analysis and display of genome-wide expression patterns. Proc Natl Acad Sci U S A 95(25): 14863–14868.

Floyd BJ, Wilkerson EM, Veling MT, Minogue CE, Xia C, Beebe ET, Wrobel RL, Cho H, Kremer LS, Alston CL et al. 2016. Mitochondrial Protein Interaction Mapping Identifies Regulators of Respiratory Chain Function. Molecular cell 63(4): 621–632.

Fruchterman TMJ, Reingold EM. 1991. Graph drawing by force-directed placement. Software: Practice and Experience 21(11): 1129–1164.

Greene CS, Krishnan A, Wong AK, Ricciotti E, Zelaya RA, Himmelstein DS, Zhang R, Hartmann BM, Zaslavsky E, Sealfon SC et al. 2015. Understanding multicellular function and disease with human tissue-specific networks. Nature genetics 47(6): 569–576.

Greene CS, Troyanskaya OG. 2012. Accurate evaluation and analysis of functional genomics data and methods. Annals of the New York Academy of Sciences 1260: 95–100.

GTEx_Consortium. 2013. The Genotype-Tissue Expression (GTEx) project. Nature genetics 45(6): 580–585.

Harrigan JA, Jacq X, Martin NM, Jackson SP. 2017. Deubiquitylating enzymes and drug discovery: emerging opportunities. Nature reviews Drug discovery.

Hein MY, Hubner NC, Poser I, Cox J, Nagaraj N, Toyoda Y, Gak IA, Weisswange I, Mansfeld J, Buchholz F et al. 2015. A human interactome in three quantitative dimensions organized by stoichiometries and abundances. Cell 163(3): 712–723.

Horlbeck MA, Xu A, Wang M, Bennett NK, Park CY, Bogdanoff D, Adamson B, Chow ED, Kampmann M, Peterson TR et al. 2018. Mapping the Genetic Landscape of Human Cells. Cell 174(4): 953–967 e922.

Huttlin EL, Bruckner RJ, Paulo JA, Cannon JR, Ting L, Baltier K, Colby G, Gebreab F, Gygi MP, Parzen H et al. 2017. Architecture of the human interactome defines protein communities and disease networks. Nature 545(7655): 505–509.

Jiang Y Oron TR Clark WT Bankapur AR D’Andrea D Lepore R Funk CS Kahanda I Verspoor KM Ben-Hur A et al. 2016. An expanded evaluation of protein function prediction methods shows an improvement in accuracy. Genome biology 17(1): 184.

Kanehisa M, Goto S, Sato Y, Furumichi M, Tanabe M. 2012. KEGG for integration and interpretation of large-scale molecular data sets. Nucleic acids research 40(Database issue): D109–114.

Khatri P, Sirota M, Butte AJ. 2012. Ten years of pathway analysis: current approaches and outstanding challenges. PLoS Comput Biol 8(2): e1002375.

Kim TH, Yoo JY, Wang Z, Lydon JP, Khatri S, Hawkins SM, Leach RE, Fazleabas AT, Young SL, Lessey BA et al. 2015. ARID1A Is Essential for Endometrial Function during Early Pregnancy. PLoS genetics 11(9): e1005537.

Klootwijk ED, Reichold M, Helip-Wooley A, Tolaymat A, Broeker C, Robinette SL, Reinders J, Peindl D, Renner K, Eberhart K et al. 2014. Mistargeting of peroxisomal EHHADH and inherited renal Fanconi’s syndrome. The New England journal of medicine 370(2): 129–138.

Kolesnikov N, Hastings E, Keays M, Melnichuk O, Tang YA, Williams E, Dylag M, Kurbatova N, Brandizi M, Burdett T et al. 2015. ArrayExpress update--simplifying data submissions. Nucleic acids research 43(Database issue): D1113–1116.

Lachmann A, Torre D, Keenan AB, Jagodnik KM, Lee HJ, Wang L, Silverstein MC, Ma’ayan A. 2018. Massive mining of publicly available RNA-seq data from human and mouse. Nature communications 9(1): 1366.

Langfelder P, Horvath S. 2008. WGCNA: an R package for weighted correlation network analysis. BMC bioinformatics 9: 559.

Lefebvre-Legendre L, Vaillier J, Benabdelhak H, Velours J, Slonimski PP, di Rago JP. 2001. Identification of a nuclear gene (FMC1) required for the assembly/stability of yeast mitochondrial F(1)-ATPase in heat stress conditions. The Journal of biological chemistry 276(9): 6789–6796.

Li H, Wang X, Rukina D, Huang Q, Lin T, Sorrentino V, Zhang H, Bou Sleiman M, Arends D, McDaid A et al. 2018. An Integrated Systems Genetics and Omics Toolkit to Probe Gene Function. Cell systems 6(1): 90–102 e104.

Li Y, Agarwal P, Rajagopalan D. 2008. A global pathway crosstalk network. Bioinformatics 24(12): 1442–1447.

Li Y, Calvo SE, Gutman R, Liu JS, Mootha VK. 2014. Expansion of biological pathways based on evolutionary inference. Cell 158(1): 213–225.

Li Y, Jourdain AA, Calvo SE, Liu JS, Mootha VK. 2017. CLIC, a tool for expanding biological pathways based on co-expression across thousands of datasets. PLoS computational biology 13(7): e1005653.

Lissanu Deribe Y, Sun Y, Terranova C, Khan F, Martinez-Ledesma J, Gay J, Gao G, Mullinax RA, Khor T, Feng N et al. 2018. Mutations in the SWI/SNF complex induce a targetable dependence on oxidative phosphorylation in lung cancer. Nature medicine 24(7): 1047–1057.

Mailman MD, Feolo M, Jin Y, Kimura M, Tryka K, Bagoutdinov R, Hao L, Kiang A, Paschall J, Phan L et al. 2007. The NCBI dbGaP database of genotypes and phenotypes. Nature genetics 39(10): 1181–1186.

Marcotte EM, Pellegrini M, Ng HL, Rice DW, Yeates TO, Eisenberg D. 1999. Detecting protein function and protein-protein interactions from genome sequences. Science 285(5428): 751–753.

Mathieu B, Sebastien H, Mathieu J. 2009. Gephi: An Open Source Software for Exploring and Manipulating Networks. International AAAI Conference on Weblogs and Social Media.

Merkwirth C, Jovaisaite V, Durieux J, Matilainen O, Jordan SD, Quiros PM, Steffen KK, Williams EG, Mouchiroud L, Tronnes SU et al. 2016. Two Conserved Histone Demethylases Regulate Mitochondrial Stress-Induced Longevity. Cell 165(5): 1209–1223.

Mitchell JA, Aronson AR, Mork JG, Folk LC, Humphrey SM, Ward JM. 2003. Gene indexing: characterization and analysis of NLM’s GeneRIFs. AMIA Annual Symposium proceedings AMIA Symposium: 460–464.

Obayashi T, Kagaya Y, Aoki Y, Tadaka S, Kinoshita K. 2019. COXPRESdb v7: a gene coexpression database for 11 animal species supported by 23 coexpression platforms for technical evaluation and evolutionary inference. Nucleic acids research 47(D1): D55–D62.

Ortega Z, Lucas JJ. 2014. Ubiquitin-proteasome system involvement in Huntington’s disease. Frontiers in molecular neuroscience 7: 77.

Pandey AK, Lu L, Wang X, Homayouni R, Williams RW. 2014. Functionally enigmatic genes: a case study of the brain ignorome. PloS one 9(2): e88889.

Pellegrini M, Marcotte EM, Thompson MJ, Eisenberg D, Yeates TO. 1999. Assigning protein functions by comparative genome analysis: protein phylogenetic profiles. Proceedings of the National Academy of Sciences of the United States of America 96(8): 4285–4288.

Pinero J, Bravo A, Queralt-Rosinach N, Gutierrez-Sacristan A, Deu-Pons J, Centeno E, Garcia-Garcia J, Sanz F, Furlong LI. 2017. DisGeNET: a comprehensive platform integrating information on human disease-associated genes and variants. Nucleic acids research 45(D1): D833–D839.

Radivojac P Clark WT Oron TR Schnoes AM Wittkop T Sokolov A Graim K Funk C Verspoor K Ben-Hur A et al. 2013. A large-scale evaluation of computational protein function prediction. Nature methods 10(3): 221–227.

Rolland T, Tasan M, Charloteaux B, Pevzner SJ, Zhong Q, Sahni N, Yi S, Lemmens I, Fontanillo C, Mosca R et al. 2014. A proteome-scale map of the human interactome network. Cell 159(5): 1212–1226.

Ross JM, Olson L, Coppotelli G. 2015. Mitochondrial and Ubiquitin Proteasome System Dysfunction in Ageing and Disease: Two Sides of the Same Coin? International journal of molecular sciences 16(8): 19458–19476.

Roy A, Kucukural A, Zhang Y. 2010. I-TASSER: a unified platform for automated protein structure and function prediction. Nature protocols 5(4): 725–738.

Schadt EE, Molony C, Chudin E, Hao K, Yang X, Lum PY, Kasarskis A, Zhang B, Wang S, Suver C et al. 2008. Mapping the genetic architecture of gene expression in human liver. PLoS biology 6(5): e107.

Schriml LM, Mitraka E, Munro J, Tauber B, Schor M, Nickle L, Felix V, Jeng L, Bearer C, Lichenstein R et al. 2019. Human Disease Ontology 2018 update: classification, content and workflow expansion. Nucleic acids research 47(D1): D955–D962.

Schroeder EA, Raimundo N, Shadel GS. 2013. Epigenetic silencing mediates mitochondria stress-induced longevity. Cell metabolism 17(6): 954–964.

Sergushichev A. 2016. An algorithm for fast preranked gene set enrichment analysis using cumulative statistic calculation. bioRxiv.

Stark C, Breitkreutz BJ, Reguly T, Boucher L, Breitkreutz A, Tyers M. 2006. BioGRID: a general repository for interaction datasets. Nucleic acids research 34(Database issue): D535–539.

Stegle O, Parts L, Piipari M, Winn J, Durbin R. 2012. Using probabilistic estimation of expression residuals (PEER) to obtain increased power and interpretability of gene expression analyses. Nature protocols 7(3): 500–507.

Stoeger T, Gerlach M, Morimoto RI, Nunes Amaral LA. 2018. Large-scale investigation of the reasons why potentially important genes are ignored. PLoS biology 16(9): e2006643.

Stroud DA, Surgenor EE, Formosa LE, Reljic B, Frazier AE, Dibley MG, Osellame LD, Stait T, Beilharz TH, Thorburn DR et al. 2016. Accessory subunits are integral for assembly and function of human mitochondrial complex I. Nature 538(7623): 123–126.

Subramanian A, Tamayo P, Mootha VK, Mukherjee S, Ebert BL, Gillette MA, Paulovich A, Pomeroy SL, Golub TR, Lander ES et al. 2005. Gene set enrichment analysis: a knowledge-based approach for interpreting genome-wide expression profiles. Proceedings of the National Academy of Sciences of the United States of America 102(43): 15545–15550.

Szklarczyk D, Franceschini A, Wyder S, Forslund K, Heller D, Huerta-Cepas J, Simonovic M, Roth A, Santos A, Tsafou KP et al. 2015. STRING v10: protein-protein interaction networks, integrated over the tree of life. Nucleic acids research 43(Database issue): D447–452.

Szklarczyk R, Megchelenbrink W, Cizek P, Ledent M, Velemans G, Szklarczyk D, Huynen MA. 2016. WeGET: predicting new genes for molecular systems by weighted co-expression. Nucleic acids research 44(D1): D567–573.

Tabach Y, Billi AC, Hayes GD, Newman MA, Zuk O, Gabel H, Kamath R, Yacoby K, Chapman B, Garcia SM et al. 2013. Identification of small RNA pathway genes using patterns of phylogenetic conservation and divergence. Nature 493(7434): 694–698.

Theisen DJ, Davidson JT, Briseño CG, Gargaro M, Lauron EJ, Wang Q, Desai P, Durai V, Bagadia P, Brickner JR et al. 2018. WDFY4 is required for cross-presentation in response to viral and tumor antigens. Science 362(6415): 694–699.

Tian Y, Garcia G, Bian Q, Steffen KK, Joe L, Wolff S, Meyer BJ, Dillin A. 2016. Mitochondrial Stress Induces Chromatin Reorganization to Promote Longevity and UPR(mt). Cell 165(5): 1197–1208.

Tong AH, Lesage G, Bader GD, Ding H, Xu H, Xin X, Young J, Berriz GF, Brost RL, Chang M et al. 2004. Global mapping of the yeast genetic interaction network. Science 303(5659): 808–813.

Uhlen M, Fagerberg L, Hallstrom BM, Lindskog C, Oksvold P, Mardinoglu A, Sivertsson A, Kampf C, Sjostedt E, Asplund A et al. 2015. Proteomics. Tissue-based map of the human proteome. Science 347(6220): 1260419.

van Dam S, Craig T, de Magalhaes JP. 2015. GeneFriends: a human RNA-seq-based gene and transcript co-expression database. Nucleic acids research 43(Database issue): D1124–1132.

Vincent DB, Jean-Loup G, Renaud L, Etienne L. 2008. Fast unfolding of communities in large networks. Journal of Statistical Mechanics: Theory and Experiment 2008(10): P10008.

Warde-Farley D, Donaldson SL, Comes O, Zuberi K, Badrawi R, Chao P, Franz M, Grouios C, Kazi F, Lopes CT et al. 2010. The GeneMANIA prediction server: biological network integration for gene prioritization and predicting gene function. Nucleic acids research 38(Web Server issue): W214–220.

Williams EG, Auwerx J. 2015. The Convergence of Systems and Reductionist Approaches in Complex Trait Analysis. Cell 162(1): 23–32.

Williams EG, Wu Y, Ryu D, Kim JY, Lan J, Hasan M, Wolski W, Jha P, Halter C, Auwerx J et al. 2018. Quantifying and Localizing the Mitochondrial Proteome Across Five Tissues in A Mouse Population. Molecular & cellular proteomics: MCP.

Wu D, Smyth GK. 2012. Camera: a competitive gene set test accounting for inter-gene correlation. Nucleic acids research 40(17): e133.

Wu Y, Williams EG, Dubuis S, Mottis A, Jovaisaite V, Houten SM, Argmann CA, Faridi P, Wolski W, Kutalik Z et al. 2014. Multilayered genetic and omics dissection of mitochondrial activity in a mouse reference population. Cell 158(6): 1415–1430.

Zhu Q, Wong AK, Krishnan A, Aure MR, Tadych A, Zhang R, Corney DC, Greene CS, Bongo LA, Kristensen VN et al. 2015. Targeted exploration and analysis of large cross-platform human transcriptomic compendia. Nature methods 12(3): 211–214, 213 p following 214.

